# Acoustoluminescence in Transition Metal and Rare Earth Oxides Beyond 1800 nm for In Vivo Imaging

**DOI:** 10.64898/2025.12.03.692246

**Authors:** Puxian Xiong, Sixin Xu, Zhexi Liu, Danyang Xu, Hanze Yu, Zideng Dai, Zhisheng Wu, Liangqiong Qu, Feifei Wang

## Abstract

Acoustoluminescence (AL) is promising for molecular imaging in living tissue, but efficient acousto-optic conversion remains challenging due to the ∼10^8^ times difference in quantum energy between phonons (1 MHz ultrasound) and photons (visible or near-infrared light). Here, we report AL in transition metal oxides (TMOs) and rare earth oxides (REOs) at wavelengths beyond 1800 nm in the short-wave infrared (SWIR) or near-infrared II (NIR-II, 1000-3000 nm) window, under ultrasound excitation at power densities 100-150-fold lower than those required for sonoluminescence in liquids. High-temperature N_2_/H_2_-mixed gas reduction was demonstrated as a safe and efficient method to regulate the AL spectra and brightness of TMOs and REOs. TMOs exhibited broadband NIR-II AL emission. Intrinsic emission peaks of rare earth ions and non-conventional luminescence were observed in the AL spectra of REOs under ultrasound excitation. NIR-II AL imaging enabled twice the penetration depth of fluorescence imaging. We developed a scanning focused ultrasound AL imaging system for *in vivo* tumor imaging through the intact hindlimb, achieving acoustic resolution and penetration depths exceeding one centimeter.

## Introduction

Optical fluorescence imaging, known for its high spatiotemporal resolution, molecular specificity^1^ and sensitivity, is extensively utilized in biology and medicine. However, its spatial resolution in deep tissue imaging rapidly deteriorates with increasing depth due to light scattering. Only multiply scattered photons have a large chance of penetrating tissue to depths beyond the optical diffusion limit, generating blurred images that lack structural details of the sample^1, 2^. In contrast, ultrasound experiences less scattering in tissues compared to light, enables better imaging resolution than magnetic resonance imaging (MRI), and is safer than positron emission tomography (PET) or computed tomography (CT), making it promising for deep tissue imaging with high resolution. However, ultrasound imaging lacks molecular specificity due to interfering signals from surrounding tissues.

Ultrasound has been introduced to enhance optical imaging in deep scattering media with acoustic resolution, such as sonoluminescent tomography^3^, ultrasound-switchable fluorescence imaging^4, 5^ and digitally time-reversed ultrasound-encoded light for fluorescence imaging^6^. For imaging modalities relying on ultrasound-excited luminescence, the initial discovery of sonoluminescence in liquid dates back to the 1930s^7^, while visible acoustoluminescence (AL) was later observed in single crystals or granular media under ultrasound stimulation^8^, such as at LiNbO_3_/air and CdS/He interfaces, and in ZnS^9^. Additionally, ultrasound has been used to induce luminescence from mechanoluminescence materials, including SrAl_2_O_4_:Eu^2+^,^10^ BaSi_2_O_2_N_2_:Eu^2+^,^11^ ZnS:Ag^+^,Co^2+^,^12^ CaZnOS:Er^3+^,^13^ CaZnOS:Nd^3+^,^14^ and organic materials^15^, with peak emissions in the visible (400-700 nm) and near-infrared I (NIR-I, 700-900 nm) spectral windows. The low luminescence intensity, short emission wavelengths (ultraviolet to NIR-I), high ultrasound power thresholds and slow response speed impose limitations on these phenomena, materials, and imaging modalities for *in vivo* deep tissue imaging.

Recently, fluorescence imaging in the near-infrared II (NIR-II, 1000-3000 nm) or short-wave infrared (SWIR) window has demonstrated higher resolution and contrast for deep tissue imaging compared to traditional visible and NIR-I imaging, due to reduced light scattering and tissue autofluorescence at longer wavelengths^1, 16–22^. This has been achieved by developing various NIR-II fluorophores/probes including rare-earth nanoparticles^23–25^, quantum dots^26, 27^, and small organic molecules^28^, and different imaging modalities, such as confocal microscopy^19^, light-sheet microscopy^22^ and structured-illumination microscopy^21^. Although NIR-II imaging allows for tissue penetration up to centimeters^1^, its resolution still remains limited by light scattering when the imaging depth exceeds the optical diffusion limit. X-ray and ultrasound sources have been applied to excite NIR-II afterglow^29–31^ to overcome excitation attenuation in tissues; however, their wide-field detection mode confines the ability to achieve high resolution imaging using multiply scattered photons.

Here, we present, for the first time, the ultrasound-excited AL from transition metal oxides (TMOs) and rare earth oxides (REOs), featuring emission peaks beyond 1800 nm in the NIR-II window. Ultrasound stimulation revealed intrinsic and non-conventional luminescence, as well as acousto-optic energy transfer in REOs. A scanning focused ultrasound AL imaging system was developed for deep tissue imaging with acoustic resolution. As an example, TiO_2_ was reduced at 1000 °C in a N_2_/H_2_ (80%/20%) atmosphere for different durations to regulate AL brightness. Reduced mesoporous silica nanoparticles (MSNs) loaded with TiO_2_ and further encapsulated with Pluronic^®^ F-127 were developed to impart aqueous solubility and biocompatibility to TiO_2_ for *in vivo* imaging. Focused ultrasound was employed to trigger AL from reduced TiO_2_, achieving twice the penetration depth compared to 1500-1700 nm NIR-II (NIR-IIb) fluorescence imaging. The tumor profile was clearly resolved through the intact mouse hindlimb by scanning focused ultrasound across it, a result not achievable with conventional NIR-IIb fluorescence imaging.

## NIR-II photoluminescence and acoustoluminescence in TMOs

Carrier recombination in TMOs has the potential to generate luminescence regardless of the presence of luminescence centers with matched energy levels^32^, as evidenced by the observation of visible and NIR-I photoluminescence (PL) in various TMOs due to different mechanisms^33–35^. To explore and regulate the PL and AL emissions from TMOs in the NIR-II window, we performed a reduction treatment under a N_2_/H_2_ (80%/20%) atmosphere at 1000 °C for 30 hours. Co_3_O_4_, NiO and CuO were instead reduced at 600 °C for 30 hours to maintain their powdered forms.

Broadband NIR-II PL emissions extending beyond 1800 nm were observed from powdered TMOs upon excitation with a focused 808-nm laser. The NIR-II PL of various TMOs exhibited distinct responses to N_2_/H_2_ reduction. As shown in Figure 1a,c-f and Supplementary Figure 1, N_2_/H_2_ reduction activated NIR-II PL in TiO_2_, enhanced NIR-II PL signal in ZrO_2_, Nb_2_O_5_ and Ta_2_O_5_, and modulated the bandwidth of NIR-II PL in V_2_O_5_, Cr_2_O_3_, MnO_2_, Fe_3_O_4_, Co_3_O_4_, NiO, CuO, MoO_3,_ and RuO_2_. No obvious changes were observed in WO_3_ arising from H_2_ reduction. The weaker PL signals observed in pristine TiO_2_, ZrO_2_, Nb_2_O_5_, and Ta_2_O_5_ compared to their reduced forms could be attributed to their lower absorption at 808 nm (Supplementary Fig. 2). Most of these NIR-II PL spectra exhibited broadband smooth profiles. A peak emission at 1365 nm appeared in pristine ZrO_2_ (Supplementary Fig. 1) but disappeared after N_2_/H_2_ reduction. It should be noted that the upper detection limit of the spectrometer used in this study was ∼2100 nm, unless otherwise noted.

**Figure 1.**
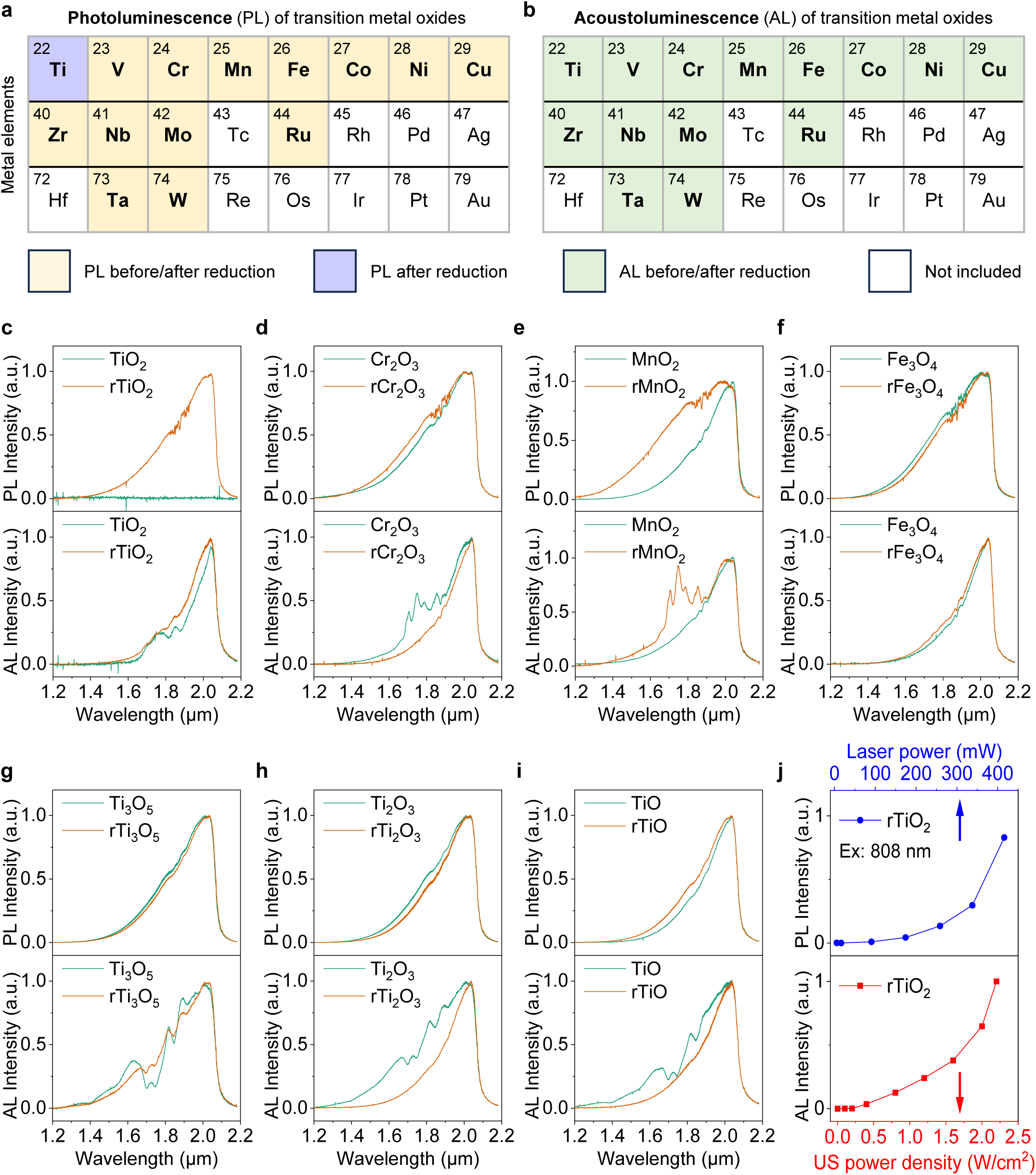
| PL and AL in TMOs beyond 1800 nm. The influence of high-temperature N_2_/H_2_ reduction on the (**a**) PL and (**b**) AL emission of TMOs. “PL before/after reduction” means that PL was observed in the TMOs both before and after N_2_/H_2_ reduction. “PL after reduction” indicates that PL was observed only in the TMOs after N_2_/H_2_ reduction. “AL before/after reduction” denotes the observation of AL in both the pristine and reduced TMOs. White-colored elements were not included in this study. We performed this reduction treatment under a N_2_/H_2_ (80%/20%) atmosphere at 1000 °C for 30 hours. (**c**-**f**) PL and AL spectra of TiO_2_, Cr_2_O_3_, MnO_2_ and Fe_3_O_4_ before and after high-temperature N_2_/H_2_ reduction. The reduced TMOs are marked with “r”. A focused 808-nm laser was used to excite the PL signal from the powdered samples. AL spectra were measured when a ∼0.5-mm layer of powdered TMOs was embedded at the bottom of a PDMS substrate and excited by an ultrasonic therapy device operating at 1 MHz. An InGaAs detector with an upper detection limit of ∼2100 nm was used to measure the spectra. The PL and AL spectra of other TMOs were summarized in Supplementary Figs. 1 and 3, respectively. (**g**-**i**) PL and AL spectra of Ti_3_O_5_, Ti_2_O_3_ and TiO before and after N_2_/H_2_ reduction, under excitation by a focused 808-nm laser and a 1 MHz ultrasonic therapy device, respectively. (**j**) Dependence curves of PL and AL intensities of rTiO_2_ on laser power and ultrasound (US) power density, respectively. These data were calculated from the spectra shown in Supplementary Fig. 6.

We recorded NIR-II AL emissions beyond 1800 nm from TMOs when a ∼0.5-mm-thick layer of powdered TMOs was embedded at the bottom of a polydimethylsiloxane (PDMS) substrate and excited by an ultrasonic therapy device operating at 1 MHz (see Methods for details). N_2_/H_2_ reduction enhanced NIR-II AL in TiO_2_, and modified the spectral profiles of NIR-II AL in Cr_2_O_3_, MnO_2_, NiO, CuO, ZrO_2_, Nb_2_O_5_, and Ta_2_O_5_ (Fig. 1b-f, Supplementary Fig. 3), whereas the spectral profiles of V_2_O_5_, Fe_3_O_4_, Co_3_O_4_, MoO_3_, RuO_2_, and WO_3_ did not change significantly. Specific emission peaks within the 1600-1900 nm window were observed in both pristine TMOs (TiO_2_, Cr_2_O_3_, ZrO_2_, Nb_2_O_5_, and Ta_2_O_5_) and reduced TMOs (MnO_2_, NiO, and CuO), while the other TMOs displayed smooth spectral profiles, similar to those observed in their PL spectra.

To explore the influence of the metal valence state, we also performed PL and AL measurements on other titanium oxides (Ti_3_O_5_, Ti_2_O_3_, and TiO; Fig. 1g-i). The broadband, smooth NIR-II PL spectra of these pristine and N_2_/H_2_-reduced titanium oxides were similar to that of N_2_/H_2_-reduced TiO_2_ (rTiO_2_). N_2_/H_2_ reduction was not essential to active their PL, which may be attributed to their higher absorption at 808 nm (Supplementary Fig. 4) and narrower bandgaps compared to TiO_2_ (Supplementary Fig. 5). NIR-II AL was observed in all titanium oxides with different spectra, and the intensities in Ti_3_O_5_, Ti_2_O_3_ and TiO were stronger than that in TiO_2_. N_2_/H_2_ reduction smoothed the AL spectral profiles of titanium oxides (Fig. 1c,g-i). Both the PL and AL intensities of rTiO_2_ increased with increasing laser power and ultrasound power density, respectively (Fig. 1j). The excitation thresholds for PL and AL of rTiO_2_ were ∼16.6 mW (focused 808-nm laser) and ∼0.1 W/cm^2^ (1 MHz ultrasound transducer), respectively (Supplementary Fig. 6).

## NIR-II photoluminescence, acoustoluminescence, and energy transfer in REOs

Rare-earth (RE) ions exhibit intrinsic luminescence arising from specific electronic transitions between internal energy levels and are widely used as activators^24, 36^. We observed both intrinsic and non-conventional broadband luminescence in REOs, induced either by laser excitation (Fig. 2a) or by acousto-optic energy transfer under ultrasound stimulation (Fig. 2b).

**Figure 2.**
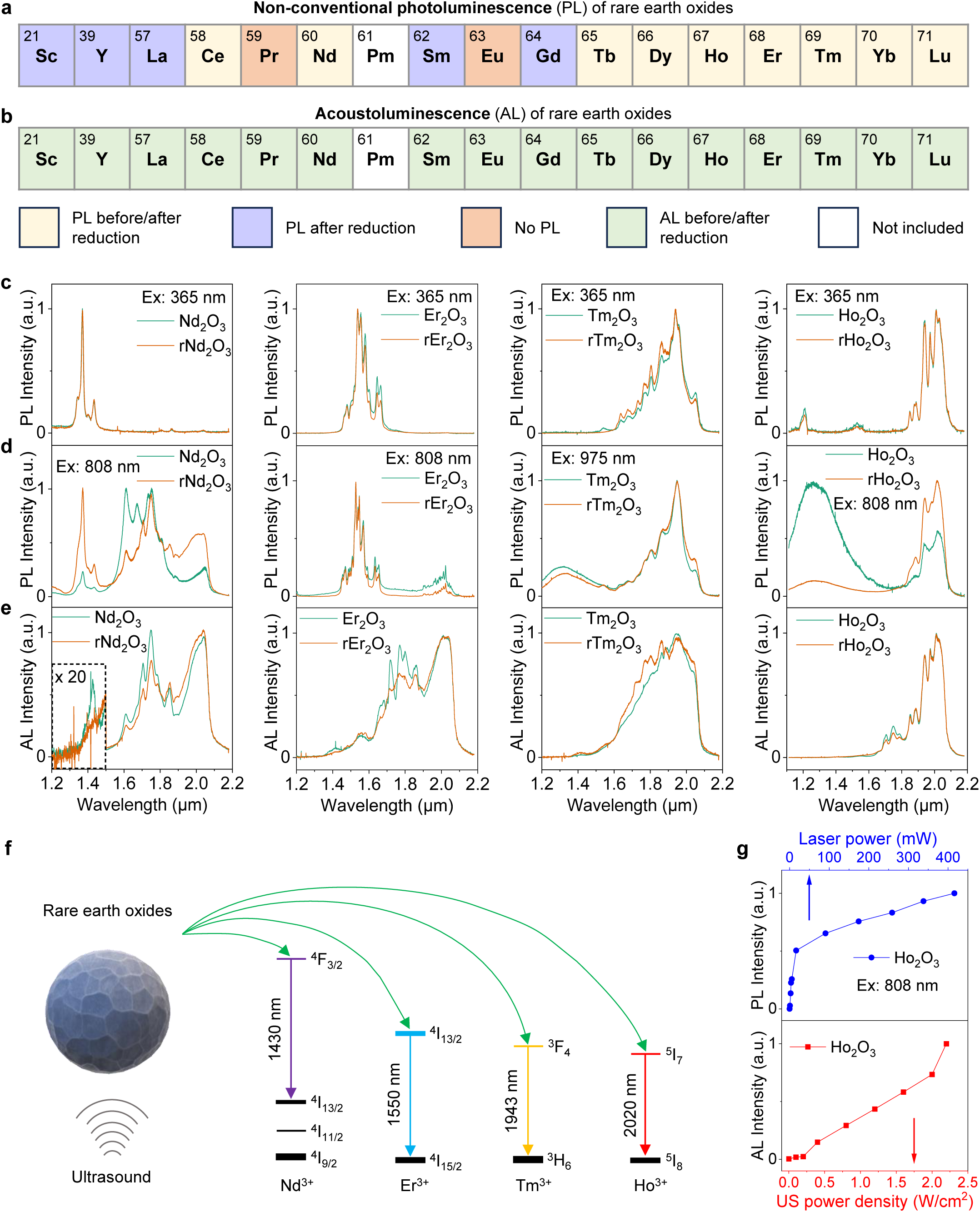
| Non-conventional PL and AL of REOs. Effects of high-temperature N_2_/H_2_ reduction on (**a**) non-conventional broadband PL and (**b**) AL emission of REOs. “No PL” indicates that no non-conventional PL was observed in the REOs before and after N_2_/H_2_ reduction. The other labels are the same as in Fig. 1. (**c**) Intrinsic PL, (**d**) non-conventional PL and (**e**) AL spectra of Nd_2_O_3_, Er_2_O_3_, Tm_2_O_3_ and Ho_2_O_3_, respectively. The AL spectra of pristine and reduced Nd_2_O_3_ between 1.2 μm and 1.5 μm were locally amplified 20 times. The intrinsic PL was excited by a 365-nm LED light. A focused 808-nm laser was used to excite the non-conventional PL signal from the powdered pristine and reduced Nd_2_O_3_, Er_2_O_3_ and Ho_2_O_3_, while a 975-nm laser was applied to pristine and reduced Tm_2_O_3_. REOs embedded in the PDMS substrate were placed on a 1 MHz ultrasonic therapy device for AL spectra measurements. The PL and AL spectra of other REOs were summarized in Supplementary Figs. 7-9. (**f**) Acousto-optic energy transfer in Nd_2_O_3_, Er_2_O_3_, Tm_2_O_3_ and Ho_2_O_3_ under ultrasound stimulation. (**g**) Influence of 808-nm laser power and US power density of a 1 MHz ultrasonic therapy device on PL and AL intensities of Ho_2_O_3_, respectively, by analyzing the spectral data shown in Supplementary Fig. 10.

Under 365-nm excitation, we observed intrinsic PL emissions (Fig. 2c) in both pristine and N_2_/H_2_-reduced Nd_2_O_3_ (Nd^3+^, ∼1370 nm, ^4^F_3/2_→^4^I_13/2_; ∼1430 nm, ^4^F_3/2_→^4^I_13/2_), Er_2_O_3_ (Er^3+^, ∼1550 nm, ^4^I_13/2_→^4^I_15/2_), Tm_2_O_3_ (Tm^3+^, ∼1943 nm, ^3^F_4_→^3^H_6_), and Ho_2_O_3_ (Ho^3+^, ∼1213 nm, ^5^I_6_→^5^I_8_; ∼2020 nm, ^5^I_7_→^5^I_8_). However, when excited with a focused 808-nm or 975-nm laser, we recorded non-conventional PL (Fig. 2d) in both pristine and N_2_/H_2_-reduced Nd_2_O_3_ (1500-2100 nm), Er_2_O_3_ (1850-2100 nm), Tm_2_O_3_ (1200-1600 nm), and Ho_2_O_3_ (1200-1750 nm), in addition to their intrinsic PL emissions. N_2_/H_2_ reduction can effectively modulate energy transfer between intrinsic and non-conventional PL in Nd_2_O_3_ and Ho_2_O_3_, as well as activate and tune broadband PL emissions in other REOs (Fig. 2a and Supplementary Fig.7).

We also detected AL in both pristine and N_2_/H_2_-reduced REOs (Fig. 2b,e, and Supplementary Fig. 8), similar to TMOs but with different spectral profiles. For REOs (Sc_2_O_3_, Y_2_O_3_, La_2_O_3_, CeO_2_, Pr_6_O_11_, Sm_2_O_3_, Eu_2_O_3_, Gd_2_O_3_, Tb_4_O_7_, Dy_2_O_3_, Yb_2_O_3_, and Lu_2_O_3_) that did not exhibit obvious intrinsic PL emission peaks in the 1400-2100 nm range (Supplementary Fig. 9), broadband AL appeared in the 1600-2100 nm range (Supplementary Fig. 8). However, the AL spectra of Nd_2_O_3_, Er_2_O_3_, Tm_2_O_3_ and Ho_2_O_3_ featured emission peaks identical to those in the intrinsic PL spectra (Fig. 2c,e). For both pristine and N_2_/H_2_-reduced Tm_2_O_3_ and Ho_2_O_3_, AL was predominantly confined to the intrinsic emission range (1600-2100 nm), while only weak intrinsic emission was present in both pristine and reduced Er_2_O_3_ at ∼1550 nm, as well as in pristine Nd_2_O_3_ at ∼1430 nm. Nd_2_O_3_ and Ho_2_O_3_ did not show intrinsic AL emission at ∼1370 nm and ∼1270 nm, respectively. The decrease in intrinsic AL emission intensity at shorter peak wavelengths could be attributed to the degree of energy matching between ultrasound-excited carriers and the intrinsic energy levels of RE ions. The RE ions can serve as illumination centers for acousto-optic energy translation for AL imaging (Fig. 2f).

The effects of laser and ultrasound power on the PL and AL of REOs were investigated, with Ho_2_O_3_ serving as an example. Both the PL and AL intensities of Ho_2_O_3_ increased with increasing laser power and ultrasound power density, with excitation thresholds of ∼0.9 mW (focused 808-nm laser) and 0.1 W/cm^2^ (1 MHz ultrasound transducer), respectively (Fig. 2g, Supplementary Fig. 10).

## Characterization of N_2_/H_2_-reduced TiO_2_

TiO_2_ is widely used in biomedical applications due to its biocompatibility and chemical stability. As an example, we modulated the luminescence of rutile TiO_2_ by reducing anatase TiO_2_ under a N_2_/H_2_ (80%/20%) gas mixture at 1000 °C, facilitating effective control of defect distribution and type in the resulting rutile TiO_2_ by adjusting the reduction time. It differs from previous methods, which produce reduced TiO_2_ in the anatase phase^37^.

As the reduction time increased, the color of TiO_2_ gradually changed from white to black (Fig. 3a and Supplementary Fig. 11), and the grain size of rTiO_2_ increased (Supplementary Fig. 12). Before reduction, the pristine TiO_2_ nanocrystals had a lattice spacing of ∼3.73 Å in the high-resolution TEM (HRTEM) image, corresponding to the (1 0 1) crystal plane of anatase TiO_2_ (Fig. 3b, left)^38^. After 30 hours of reduction, the grains grew to several micrometers, with a lattice spacing of ∼2.56 Å corresponding to the (2 0 16) crystal plane of Ti_9_O_17_ (Fig. 3b, right)^39^.

**Figure 3.**
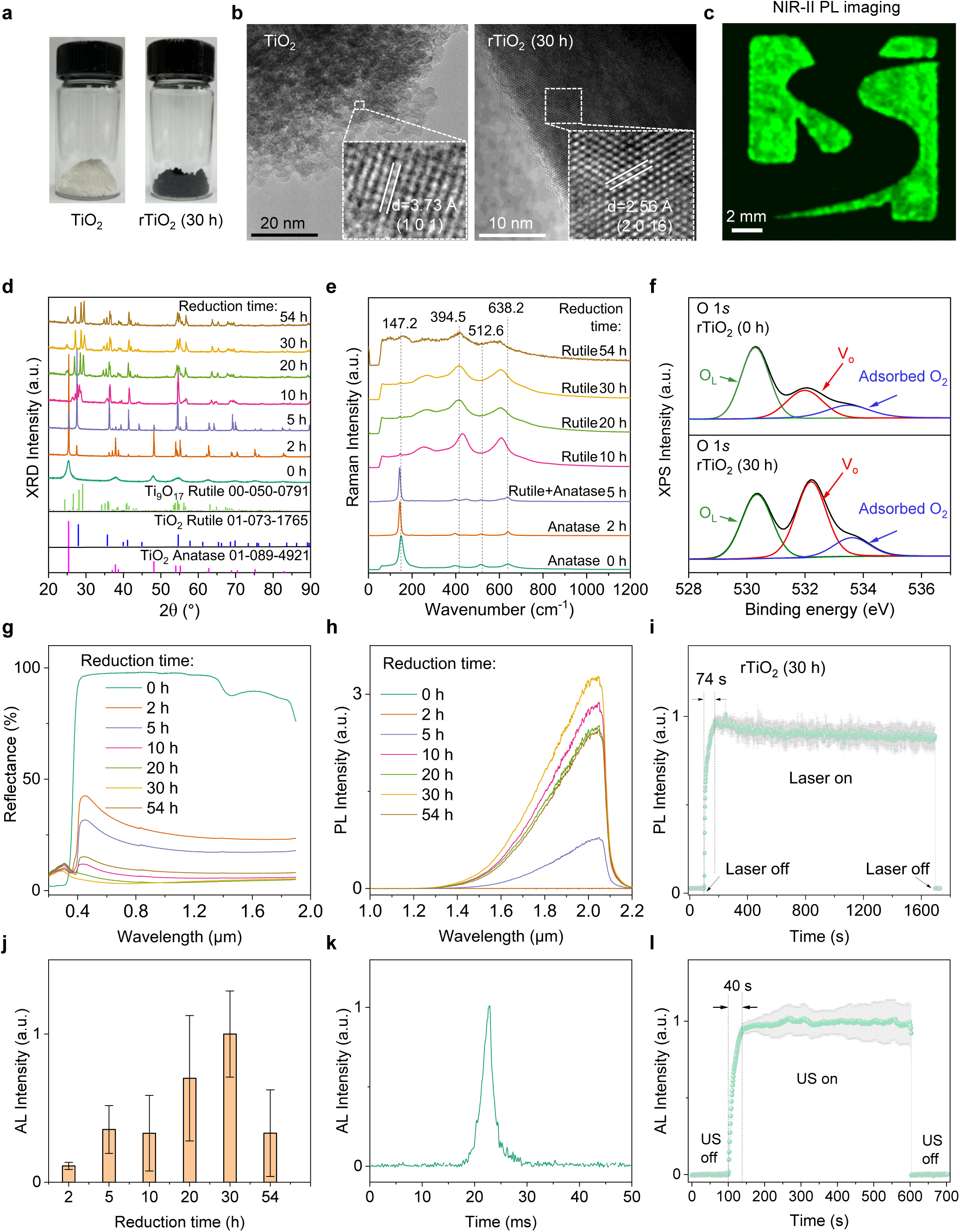
| Characterization and optimization of NIR-II PL and AL of rTiO_2_. (**a**) Photographs show the color changes of TiO_2_ before and after 30 hours of N_2_/H_2_ reduction. (**b**) High-resolution TEM images illustrate the lattice changes before and after N_2_/H_2_ reduction. (**c**) Scanning NIR-II PL imaging of rTiO_2_-embedded PDMS substrate in the shape of the ’MILES’ logo. The sample was scanned across a focused 660-nm laser. The PL was collected by an InGaAs PMT after passing through a 1000-nm long-pass filter. The raster-scanning step size and speed of the motorized stage were 0.1 mm and 5 mm/s, respectively. (**d**) XRD patterns, (**e**) Raman spectra, (**f**) O 1*s* XPS spectra, (**g**) diffuse reflectance spectra, and (**h**) PL spectra under focused 660-nm laser excitation of TiO_2_ before and after 2, 5, 10, 20, 30 or 54 hours of N_2_/H_2_ reduction at 1000 °C. (**i**) PL stability of rTiO_2_ reduced for 30 hours. A 660-nm laser was used for excitation. (**j**) The influence of N_2_/H_2_ reduction time on the AL intensity of rTiO_2_. A ∼0.5-mm-thick layer of powdered rTiO_2_ was embedded at the bottom of a PDMS substrate and positioned at the focus of an ultrasound transducer operating at ∼4.55 MHz for AL measurements. The AL emission was collected by an InGaAs camera in the 1000-1700 nm window. (**k**) Transient response of AL emission from rTiO_2_ reduced for 30 hours under pulsed ultrasound with a duration of 10 ms. (**l**) AL stability of rTiO_2_ reduced for 30 hours. In (**i**), (**j**) and (**l**), data are shown as mean ± s.d. derived from analyzing 3, 25 and 8 measured values, respectively.

As shown in the X-ray diffraction (XRD) results (Fig. 3d), the broader XRD peaks indicated weaker crystallinity and smaller grain size in pristine anatase TiO_2_ than in rTiO_2_. Extending the reduction time gradually transformed TiO_2_ from its initial anatase phase at 0 hours to a mixture of anatase and rutile phases at 2 and 5 hours. Sharp XRD diffraction peaks were observed, indicating a highly crystallized phase after high-temperature N_2_/H_2_ reduction. After 10 hours of reduction, only the rutile phase was observed, eventually leading to the formation of Ti_9_O_17_ rich in oxygen vacancies, in agreement with HRTEM observations (Fig. 3b).

The structural changes in TiO_2_ induced by N_2_/H_2_ reduction can be further assessed by Raman spectroscopy. Four Raman peaks at 147.2, 394.5, 512.6, and 638.2 cm^-1^, corresponding to the anatase phase of TiO_2_,^40^ were observed before reduction (Fig. 3e). As the reduction time was extended to 5 hours, a weak Raman peak at 424.7 cm^-1^ emerged. Ten hours after reduction, rutile TiO_2_’s Raman peaks at 247.8, 424.7, and 608.9 cm^-1^ were resolved while the typical anatase Raman bands disappeared, confirming the phase change from anatase to rutile^41^.

High-temperature N_2_/H_2_ is expected to create abundant defects on the surface or subsurface of TiO_2_^.42^ We assessed the changes in surface chemical bonding of TiO_2_ induced by high-temperature N_2_/H_2_ reduction for various durations using X-ray photoelectron spectroscopy (XPS).

The O 1*s* XPS spectra showed distinct peaks at 530.3 eV and 532.2 eV, corresponding to O species in Ti-O and surface Ti-OH, respectively (Supplementary Fig. 13a). The O 1*s* XPS spectra can be deconvoluted into three Gaussian components (Fig. 3f), assigned to lattice oxygen (O_L_), oxygen vacancy (V_o_) and adsorbed O_2_^42^. N_2_/H_2_ reduction for 30 hours can greatly increase the component of V_o_ in rTiO_2_. The Ti 2*p* XPS peaks at 459.1 eV and 465.1 eV, corresponding to Ti 2*p*_3/2_ and 2*p*_1/2_ peaks, were observed in all samples (Supplementary Fig. 13b). Further deconvolution of Ti 2*p* XPS spectra revealed that N_2_/H_2_ reduction increases the component of low-valence titanium ions in rTiO_2_, consistent with the introduction of V_o_ after reduction (Supplementary Fig. 13c). The hydroxyl group was formed on the surface of rTiO_2_^,42^ while V_o_ may be present in the subsurface, as indicated by faint electron paramagnetic resonance (EPR) signals measured at room temperature (Supplementary Fig. 14a). Low-temperature (100 K) EPR signals of rTiO_2_ reduced for 0 hours and 30 hours further confirmed the existence of Ti^3+^-V_o_ associates (Supplementary Fig. 14b).

We examined the diffuse reflectance spectra of TiO_2_ reduced for different durations (Fig. 3g). Both the TiO_2_ before and after reduction had strong absorption in the ultraviolet spectra band (< 420 nm). The pristine anatase TiO_2_ exhibited strong reflectance at wavelengths greater than 420 nm. Upon reduction, the reflectivity of TiO_2_ significantly decreased across the 400-1900 nm range, particularly after 20 or 30 hours of reduction. This enhanced visible and near-infrared absorption could be attributed to the creation of V_o_ in rTiO_2_ during high-temperature N_2_/H_2_ treatment^42^, which reshaped the bandgap structure^43^. The bandgaps of rTiO_2_ samples reduced for 0, 2, 5, 10, 20, 30, and 54 hours were 3.66, 3.21, 3.00, 2.87, 0.80, 0.73, and 2.89 eV, respectively (Supplementary Fig. 15).

## Optimization of NIR-II photoluminescence and acoustoluminescence in N_2_/H_2_-reduced TiO_2_

The pristine TiO_2_ (0 hours) and lightly reduced TiO_2_ (2 hours) exhibited no obvious NIR-II PL emissions (Fig. 3h). The NIR-II PL intensity varied with reduction time and temperature, reaching a maximum at 30 hours and 1000 °C (Supplementary Fig. 16). An emission peak at ∼2280 nm was measured using an indium gallium arsenide (InGaAs)-based spectrometer with an upper detection limit of 2500 nm (Supplementary Fig. 17), which differs from the previously reported visible and NIR-I PL of anatase and rutile TiO_2_.^33, 44^ The spectral profiles remained unchanged when we excited rTiO_2_ using lasers in the 660-1020 nm range (Supplementary Fig. 18). The PL signal responded rapidly to 660-nm laser pulses with durations ranging from 1 to 100 ms (Supplementary Fig. 19). After the 660-nm laser was turned on, the PL signal increased during the initial ∼74 seconds and then stabilized at a high intensity level over ∼30 minutes (Fig. 3i). No persistent luminescence was observed when the laser was switched off.

We performed NIR-II PL imaging of rTiO_2_ embedded at the bottom of a PDMS substrate in the shape of the ’MILES’ logo with a thickness of ∼0.5 mm. The sample was placed on a motorized scanning stage and excited with a focused 660-nm laser. The PL was collected in the 1000-1700 nm window. NIR-II PL imaging was achieved by raster-scanning the sample across the focused laser (Fig. 3c).

Similar to the brightest PL signal (Fig. 3h), the strongest AL signal was also observed in the 30-hour rTiO_2_ (Fig. 3j, see Methods for details). The AL signal responded quickly to ultrasound with pulse widths ranging from 1 to 100 ms (Fig. 3k and Supplementary Fig. 20). As the ultrasound was turned on, the AL in rTiO_2_ increased during the initial 40 seconds and subsequently stabilized over a period of ∼8 minutes (Fig. 3l). No afterglow signal was recorded after the ultrasound transducer was turned off.

## Phantom and *in vivo* NIR-II acoustoluminescence imaging

The average grain size of directly reduced TiO_2_ at 1000 °C for 30 hours was ∼3.75 µm (Supplementary Fig. 21), making it incompatible with biomedical applications. To decrease the grain size, we employed an MSN-based template for the reduction of TiO_2_ (rTiO_2_-MSN; Methods), achieving a stronger AL signal than rTiO_2_ (Fig. 4a). The brightest AL of rTiO_2_-MSN was achieved by reduction at 1000 °C for 30 hours with a mass ratio of MSN to TiO_2_ of 2:1 (Fig. 4b and Supplementary Fig. 22). TEM results revealed MSN was partially coated on the surface of rTiO_2_ (Supplementary Fig. 23). Pluronic^®^ F-127 was used to encapsulate rTiO_2_-MSN (rTiO_2_-MSN@F127) to impart aqueous solubility and biocompatibility, and to minimize water-induced quenching of luminescence (Fig. 4c). The average hydrated size of rTiO_2_-MSN@F127 was measured by dynamic light scattering (DLS) to be ∼420 nm (Fig. 4d).

**Figure 4.**
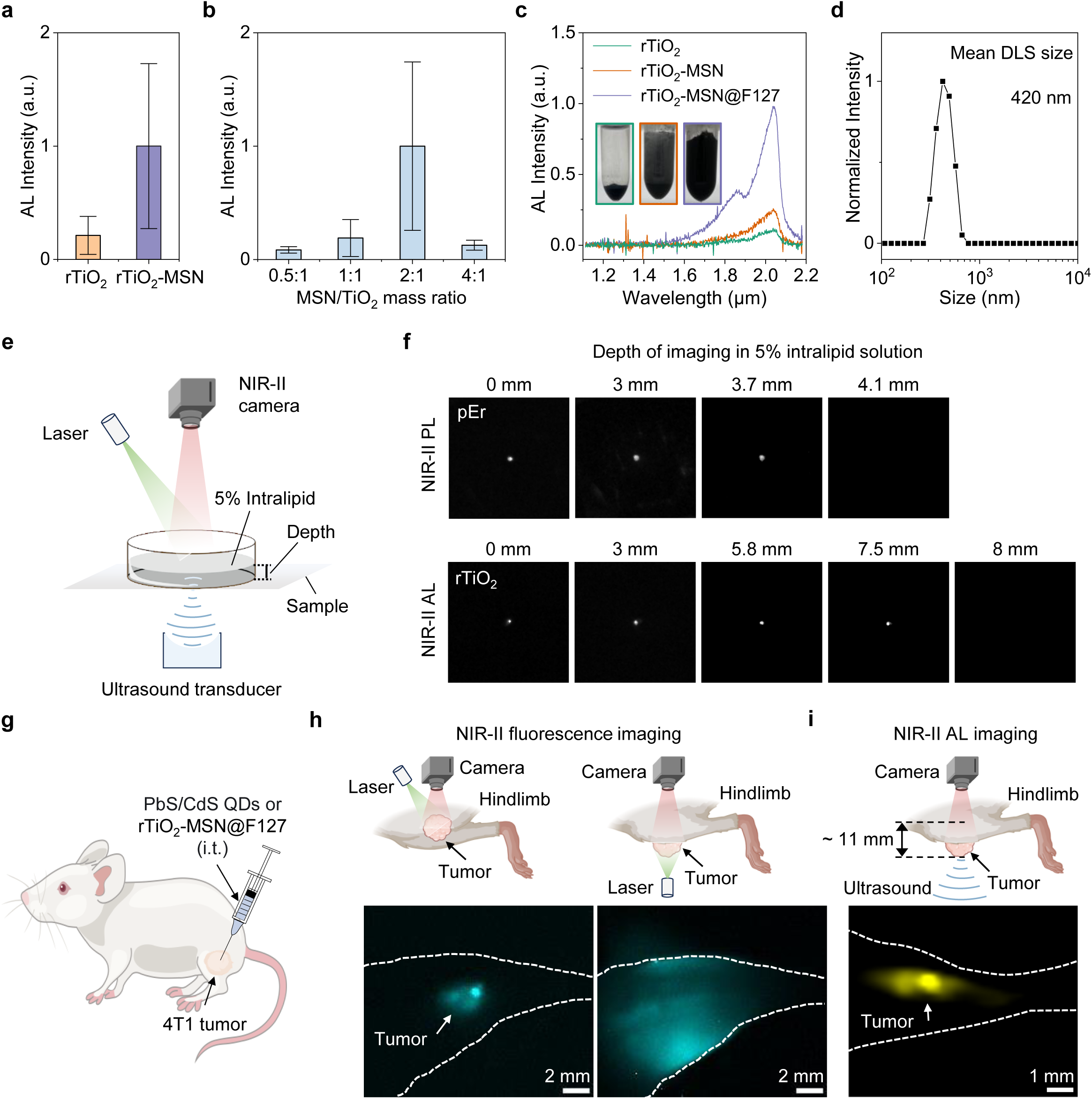
| *In vivo* NIR-II AL tumor imaging. (**a**) Comparison of AL intensity between rTiO_2_ and TiO_2_ reduced in MSN-based template (rTiO_2_-MSN). High-temperature reduction was performed in a N_2_/H_2_ (80%/20%) gas mixture at 1000 °C for 30 hours. (**b**) AL intensity of rTiO_2_-MSN with different MSN/TiO_2_ mass ratios, reduced at 1000 °C for 30 hours in a N_2_/H_2_ (80%/20%) gas mixture. (**c**) AL spectra of rTiO_2_, rTiO_2_-MSN and rTiO_2_-MSN@F127 in PBS solution, with corresponding photographs shown in the inset. Pluronic^®^ F127 encapsulation imparted aqueous solubility and biocompatibility to rTiO_2_-MSN. (**d**) Dynamic light scattering spectra of rTiO_2_-MSN@F127 in PBS buffer. (**e**) A simplified schematic of the imaging system for comparing NIR-II fluorescence imaging and scanning focused ultrasound AL imaging. A layer of 5% intralipid solution with different thicknesses was used for phantom imaging to mimic brain tissue scattering. (**f**) Comparison of penetration depths of NIR-IIb fluorescence imaging and NIR-II AL imaging. For fluorescence imaging, a drop of pEr (emission peak: ∼1550 nm)^25^ was placed beneath a layer of 5% intralipid solution with different thicknesses. A 975-nm laser was directed at an angle, passed through the intralipid and excited the pEr. Fluorescence in the 1500-1700 nm window was collected by an InGaAs camera positioned above the intralipid. In ultrasound-triggered NIR-II AL imaging, a rTiO_2_-embedded PDMS sample was positioned at the focus of an ultrasound transducer operating at ∼4.55 MHz for acoustic excitation. The AL emission was collected by the same camera after passing through a layer of 5% intralipid solution. (**g**) Intratumoral injection of PbS/CdS QDs (peak emission: ∼1600 nm)^26^ or rTiO_2_-MSN@F127 for *in vivo* fluorescence imaging and scanning focused ultrasound AL imaging. A 4T1 breast tumor was inoculated on the hindlimb of BALB/c mice and probes were administered intratumorally when the tumor size reached ∼4 × 4 × 2.8 mm^3^ (length × width × thickness). (**h**) *In vivo* NIR-II fluorescence imaging of a tumor positioned facing the NIR-II camera (left) or placed beneath the hindlimb (right). An 808-nm laser was used for excitation and fluorescence was collected in the 1500-1700 nm. Strong scattering by the hindlimb limits tumor profiling through the hindlimb using fluorescence imaging. (**i**) Scanning focused ultrasound NIR-II AL imaging of a tumor beneath the hindlimb, facilitating tumor profiling with acoustic resolution and a penetration depth of ∼11 mm. An ultrasound transducer operating at ∼4.55 MHz was used for acoustic excitation. AL was collected by an InGaAs camera synchronized with a motorized stage. The exposure time of the camera was 90 ms. The motorized stage was raster-scanned with a step size of 0.2 mm and a speed of 1.5 mm/s. In (**a**) and (**b**), data are shown as mean ± s.d., derived from analyzing ∼55 and ∼107 measured values, respectively.

We performed phantom imaging to compare the imaging depth of laser-excited NIR-IIb fluorescence imaging with ultrasound-triggered NIR-IIb AL imaging. For fluorescence imaging, a drop of NaErF_4_/NaYF_4_ core-shell down-conversion nanoparticles^25^ (pEr, emission peak: ∼1550 nm) was placed beneath a layer of 5% intralipid solution with different thicknesses to mimic brain tissue scattering^1^. A 975-nm laser was directed at an angle, passed through the intralipid and excited the pEr. An InGaAs camera positioned above the intralipid was used to collect the fluorescence signal after it was filtered by a 1500-nm long-pass filter (Fig. 4e). The penetration limit of NIR-IIb fluorescence imaging was ∼3.7 mm (Fig. 4f, top), consistent with previous observations^19^. For ultrasound-triggered NIR-IIb AL imaging, a rTiO_2_-embedded PDMS sample was positioned at the focus of an ultrasound transducer operating at ∼4.55 MHz, and the AL was collected by the same camera after passing through a layer of 5% intralipid solution (Fig. 4e). NIR-IIb AL imaging enabled a penetration depth of ∼7.5 mm (Fig. 4f, bottom), about twice that of fluorescence imaging. In fluorescence imaging, both excitation and emission light are attenuated as they penetrate. Since ultrasound is scattered and attenuated less than light, ultrasound-excited AL imaging enables deeper tissue penetration.

To evaluate the differences between NIR-II fluorescence and AL imaging *in vivo*, we performed tumor imaging in live mice. A 4T1 breast tumor was inoculated on the hindlimb of BALB/c mice. We intratumorally (i.t.) injected either core/shell lead sulfide/cadmium sulfide quantum dots (PbS/CdS QDs, peak emission: ∼1600 nm)^26^ or rTiO_2_-MSN@F127 into the tumor when its size reached ∼4 × 4 × 2.8 mm^3^ (length × width × thickness; Fig. 4g). For NIR-IIb fluorescence imaging, an 808-nm laser was used to excite PbS/CdS QDs, and the emitted fluorescence was collected in the 1500-1700 nm window. As the tumor was positioned facing the NIR-II wide-field imaging system, the boundary of the PbS/CdS QDs-labeled tumor could be clearly resolved (Fig. 4h, left). However, when the tumor was placed beneath the mouse hindlimb with a thickness of ∼8.28 ± 0.26 mm and imaged through the hindlimb, only scattered fluorescence signals without tumor profile information were detected (Fig. 4h, right).

For NIR-IIb AL imaging, we developed a scanning focused ultrasound AL imaging system for *in vivo* imaging (Supplementary Fig. 24, see Methods for details). We performed ultrasound-triggered NIR-II AL imaging by scanning the tumor across the focus of an ultrasound transducer, which had a frequency of ∼4.55 MHz and a full width at half maximum (FWHM) of ∼1 mm. (Supplementary Fig. 25). The AL signal was collected by an extended InGaAs detector with an upper detection wavelength limit of ∼1900 nm. In contrast to fluorescence imaging (Fig. 4h, right), the tumor edge could be clearly resolved with acoustic resolution when the AL signal was collected after transmission through the intact mouse hindlimb, which comprised skin, bone, and muscle (Fig. 4i). The total thickness of hindlimb and tumor was ∼11 mm, indicating that our method enables high-resolution, centimeter-depth live tissue imaging.

## Discussion

Ultrasound-excited luminescence relies on acousto-optic conversion, with its efficiency being crucial for achieving deep penetration in live tissue imaging. Sonoluminescence and ultrasound-luminescence conversion materials (ULCM) usually require a high ultrasound power density (sonoluminescence: 10-15 W/cm^2^;^45^ NIR-I ULCM: ∼1.4-1117.4 W/cm^2^, Supplementary Table 1), long excitation time (seconds to minutes)^15^ and exposure time (seconds to minutes)^46^ for *in vivo* imaging, due to the low energy conversion efficiency. Their visible and NIR-I luminescence, combined with the wide-field detection modality, still limits the capability for high-resolution deep tissue imaging because of strong scattering at short wavelengths, although ultrasound-excited luminescence imaging facilitates deeper penetration than fluorescence imaging due to less scattering experienced by ultrasound than light excitation.

In this work, we explored NIR-II acoustoluminescence in TMOs and REOs and observed luminescence beyond 1800 nm under ultrasound excitation at a power density of ∼0.1 W/cm^2^ (Supplementary Figs. 6 and 10), which was 14-11174 times lower than that required by previous materials. The underlying mechanism of AL generation in TMOs and REOs still lacks comprehensive understanding. TMOs and REOs exhibited different AL emission properties. In TMOs, AL emission may arise from radiative transitions between donor and acceptor levels, following their recharging via acoustically stimulated charge carriers^9, 47^. In REOs, we observed intrinsic emission peaks under ultrasound excitation, revealing acousto-optic energy transfer to RE ions. RE ions can serve as activators for AL imaging if the energy of ultrasound-excited charge carriers matches their intrinsic energy levels. High-temperature N_2_/H_2_-mixed gas reduction provided a safe and effective approach for modulating the spectral profile and intensity in TMOs and REOs, indicating the potential influence of defects on AL emission.

AL imaging enabled twice the penetration depth of fluorescence imaging and had the capacity to extend imaging depth well beyond the optical diffusion limit, while maintaining acoustic resolution. By developing a scanning focused ultrasound AL imaging system (Supplementary Fig. 24), we mapped tumor profiles *in vivo* through the intact mouse hindlimb with acoustic resolution and a penetration depth of ∼11 mm, whereas they were completely obscured in NIR-IIb fluorescence imaging. AL imaging allows for high molecular specificity, which is difficult to achieve with ultrasound imaging and photoacoustic imaging due to their inevitable background signals from tissues. Compressed sensing methods can be employed to compensate for the slow speed of raster scanning. The operating frequency (∼4.55 MHz) of the focused ultrasound transducer confines the AL imaging resolution to ∼1 mm, which can be improved by using transducers with higher frequencies. For example, the lateral FWHM at the focus of a 50 MHz focused ultrasound transducer is ∼50-100 µm. Super-resolution artificial intelligence networks can also be introduced to further enhance resolution. Advances in AL imaging modalities and materials will open up new opportunities for biomedical imaging with high resolution, molecular sensitivity and penetration depth beyond optical diffusion limit.

## Methods Materials

TiO_2_ (99.8%, metals basis, 5-10 nm, anatase), V_2_O_5_ (99.99%, metals basis), Cr_2_O_3_ (99.95%, metals basis), MnO_2_ (99.95%, metals basis), Fe_3_O_4_ (≥ 99%, metals basis, powder, 20 nm), Co_3_O_4_ (≥ 99.99%, trace metals basis), NiO (99.99%, metals basis), CuO (99.9%, metals basis), ZrO_2_ (99.99%, metals basis, Removal of Hf or HfO_2_), Nb_2_O_5_ (99.99%, metals basis), MoO_3_ (99.9%, metals basis), RuO_2_ (99.9%, metals basis), Ta_2_O_5_ (99.99%, metals basis), WO_3_ (99.8%, metals basis), Sc_2_O_3_ (≥ 99.9%, metal basis), Y_2_O_3_ (≥ 99.9%, metal basis), La_2_O_3_ (≥ 99.99%, metal basis), CeO_2_ (≥ 99.95%, metal basis, white powder), Pr_6_O_11_ (≥ 99.9%, metal basis), Nd_2_O_3_ (≥ 99.9%, metal basis), Sm_2_O_3_ (≥ 99.9%, metal basis), Eu_2_O_3_ (≥ 99.9%, metal basis), Gd_2_O_3_ (≥ 99.9%, metal basis), Tb_4_O_7_ (≥ 99.9%, metal basis), Dy_2_O_3_ (≥ 99.9%, metal basis), Ho_2_O_3_ (≥ 99.9%, metal basis, < 100 nm particle size (SEM)), Er_2_O_3_ (≥ 99.9%, metal basis), Tm_2_O_3_ (≥ 99.9%, metal basis), Yb_2_O_3_ (≥ 99.99%, metal basis), Lu_2_O_3_ (≥ 99.9%, metal basis), and SiO_2_ (mesostructured, MCM-41 type (hexagonal)) were purchased from Aladdin without further purification. Pluronic^®^ F-127 (C_3_H_6_O·C_2_H_4_O)_x_ was obtained from Sigma-Aldrich. SYLGARD^TM^ 184 Silicone Elastomer was purchased from Dow Corning. Ultrasound gel (Kefu D-ⅡⅩ) was purchased from Ningjin County Jinyang Medical Materials Factory.

## Preparation of N_2_/H_2_-reduced TMOs and REOs

Reduced TMOs and REOs were obtained by directly reducing powdered raw materials in a horizontal tube furnace (OTF-1200X, HF-Kejing) under a flowing N_2_/H_2_ (80%/20%)-mixed gas environment (flow rate: ∼4.44 cc/min) at 1000 °C for 30 hours. For each reduction procedure, the temperature ramp rate was 5 °C/min from 25 to 800 °C, and 3 °C/min from 800 to 1000 °C. The cooling rate was ∼1.4 °C/min. To preserve the powdered form of the target samples, Co_3_O_4_, NiO and CuO were reduced at 600 °C. For these three materials, the temperature ramp rate was 5 °C/min from 25 to 600 °C. The cooling rate was ∼1.4 °C/min.

## AL and PL spectra measurements

Both AL and PL spectra were measured using a spectrometer (Spectra Pro HRS-300-MS, Teledyne Vision Solutions) equipped with a liquid-nitrogen-cooled InGaAs linear camera with an upper detection limit of ∼2100 nm (PyLoN-IR 2.2, Teledyne Vision Solutions).

To measure the AL spectra, a ∼0.5-mm-thick layer of powdered TMOs and REOs was embedded at the bottom of a PDMS substrate (TMOs/REOs-PDMS) and positioned on the working plane of an ultrasonic therapy device (Sonicator 740, Mettler Electronics) operating at 1 MHz. The material surface was positioned directly facing the detection system to prevent interference from PDMS absorption, and a layer of 0.5-1 cm ultrasound gel was applied between the ultrasonic therapy device and TMOs/REOs-PDMS to match the acoustic impedance.

To measure the non-conventional broadband PL spectra, pristine and reduced TMOs and REOs powders were placed on weighing papers and positioned at the focal point of the focused laser with a wavelength in the range of 660-1020 nm. Intrinsic spectra of powdered REOs were excited by a 365-nm UV LED.

## Synthesis of rTiO_2_

To modulate the AL and PL performances of rTiO_2_, we adjusted both the reduction temperature (200, 400, 600, 800, and 1000 °C) and reduction time (0, 2, 5, 10, 20, 30, and 54 hours) (Fig. 3h,j). rTiO_2_ was synthesized by reducing TiO_2_ (99.8%, metals basis, 5-10 nm, anatase) using a horizontal tube furnace (OTF-1200X, HF-Kejing) in a flowing N_2_/H_2_ (80%/20%)-mixed gas environment (flow rate: ∼4.44 cc/min). For each sintering procedure, the temperature ramp rate was 5 °C/min from 25 to 800 °C, and 3 °C/min from 800 to 1000 °C. The cooling rate was ∼1.4 °C/min.

## Synthesis of rTiO_2_-MSN nanocrystals

rTiO_2_-MSN nanocrystals were prepared by an MSN-restricted growth method. MSN and TiO_2_, with different mass ratios (0.5:1, 1:1, 2:1, and 4:1), were mixed in 10 mL of ethanol and sonicated for 10 minutes. The obtained solution was poured into an Al_2_O_3_ crucible boat and reduced in a flowing N_2_/H_2_ (80%/20%) atmosphere (flow rate: ∼4.44 cc/min) in a tube furnace (OTF-1200X, HF-Kejing) at a reduction temperature of 800, 900, 1000, 1100, or 1200 °C. The temperature ramp rate was 5 °C/min from 25 to 800 °C, and 3 °C/min from 800 to 1200 °C. The cooling rate was ∼1.4 °C/min. The obtained powders were ground for 2 minutes before further use. The reduction temperature and time (10, 20, 30, 40, and 54 hours) were optimized to achieve the brightest AL (Supplementary Fig. 22).

## Preparation of rTiO_2_-MSN@F127 PBS solution

40 mg rTiO_2_-MSN nanocrystals were added to 10 mL of chloroform and sonicated for 5 minutes. 160 mg of Pluronic^®^ F-127 was added to this solution and sonicated for 10 minutes. The resulting solution was stirred overnight until the chloroform was completely evaporated. Then, 10 mL of deionized water was added, and the mixture was sonicated for 8-10 minutes, followed by centrifugation at 13,000 rpm for 8 minutes. The precipitate was washed two times with deionized water to remove excess Pluronic^®^ F-127. The obtained rTiO_2_-MSN@F127 was dispersed in 1 mL PBS to obtain a 40 mg/mL rTiO_2_-MSN@F127 PBS solution for further *in vivo* imaging.

## Materials characterization

TEM images were captured by a FEI Talos 200x transmission electron microscope. XRD patterns of rTiO_2_ were obtained with a Bruker D8 ADVANCE powder diffractometer. Raman spectra of rTiO_2_ were recorded with a Renishaw inVia Raman spectrometer using a 25-mW Nd:YAG laser at 532 nm. XPS spectra of TiO_2_ and rTiO_2_ were collected with an ESCALAB QXi XPS microprobe. Diffuse reflectance spectra of rTiO_2_ were obtained using a UH4150 UV-VIS-NIR spectrophotometer (Hitachi), with white Al_2_O_3_ powder as a reference. SEM images were taken using a Helios 5CX scanning electron microscope equipped with an XFlash 7100 energy-dispersive spectrometer. EPR spectra of TiO_2_ and rTiO_2_ were collected with an EMXplus X-band spectrometer (Bruker, Germany) equipped with a CH-210N Cryocooler (Cold Edge). Material temperatures were measured using a thermal imager (HT-A2, Hti).

## Cell culture, tumor inoculation and mouse use for *in vivo* imaging

All animal experiments were performed under the approval of The University of Hong Kong’s Administrative Panel on Laboratory Animal Care. All experiments were performed in accordance with the Centre for Comparative Medicine Research of Laboratory Animals in Hong Kong. Six-week-old BALB/c female mice were purchased from the Centre for Comparative Medicine Research, The University of Hong Kong. The 4T1 cells were cultured in Roswell Park Memorial Institute (RPMI) 1640 medium supplemented with 10 % (v/v) Fetal Bovine Serum (FBS) at 37 °C in a 5 % (v/v) CO_2_ humidified atmosphere. 4T1 cells (1 × 10^7^ cells) suspended in 100 μL PBS were subcutaneously injected to each mouse to establish a 4T1 tumor-bearing mouse model. Mice were selected randomly from cages for experiments.

## Wide-field NIR-II fluorescence imaging

For wide-field imaging shown in Fig. 4h, PbS/CdS QDs (peak emission: ∼1600 nm)^26^ were injected intratumorally into the 4T1 tumor when the tumor size reached ∼4 × 4 × 2.8 mm^3^ (length × width × thickness). *In vivo* NIR-II fluorescence imaging of a tumor positioned facing the NIR-II camera or placed beneath the hindlimb was performed in a NIR-II small animal imaging system (KingsVision™, Nirmidas Biotech). The QDs were excited using an 808-nm laser with excitation power density of ∼100 mW/cm^2^ at the imaging plane. The fluorescence was filtered by two 1500-nm long-pass filters for NIR-IIb imaging. The exposure time of Fig. 4h (left) and Fig. 4h (right) were 0.1 ms and 15 ms, respectively.

## Scanning focused ultrasound AL imaging system

Focused ultrasound was generated using an ultrasound transducer operating at ∼4.55 MHz with a FWHM of ∼1 mm (Supplementary Fig. 25). The ultrasound transducer was driven by a custom power amplifier with a sine wave input from a function generator (SDG2122X, SIGLENT), and impedance matching for the transducer was achieved using a matching network. The ultrasound-triggered AL was collected using a fixed-focal lens (SWIR-25, Navitar) and an InGaAs camera (C-RED 2 ER, Oxford Instruments). A motorized stage (M-VP-25XL, Newport Corporation) was used for raster scanning and controlled by a LabVIEW program. The motorized stage and camera were synchronized via the stage controller (XPS-RLD4, Newport Corporation).

## *In vivo* scanning focused ultrasound AL imaging

For *in vivo* scanning focused ultrasound AL imaging, a 4T1 breast tumor was inoculated on the hindlimb of BALB/c mice. When the tumor size reached ∼4 × 4 × 2.8 mm^3^ (length × width × thickness), 50 μL of rTiO_2_-MSN@F127 PBS solution was injected intratumorally (Fig. 4g). After anesthetization, the mouse was mounted on the motorized stage with the tumor facing the ultrasound transducer (Fig. 4i). Ultrasound-triggered AL signals from the tumor penetrated through the intact hindlimb and were collected by an InGaAs camera. The scan area was 10 × 10 mm^2^, with a step size of 0.2 mm. The scanning speed was 1.5 mm/s. The exposure time of the InGaAs camera was 90 ms, and a 1500-nm long-pass filter was used to filter the AL emission. The acquired multi-frame image sequence was used to reconstruct an image in MATLAB.

## Statistical and data analysis

Data analysis was performed using MATLAB (2022) or OriginLab (2024b). Nano Measure (Version 1.2.5) was used to calculate the average grain size. The integrated NIR-II PL/AL intensity values shown in Figure 1j and Figure 2g, the average PL or AL intensity values and standard deviations presented in Figure 3i,j,l, Figure 4a,b, Supplementary Figure 21b, and Supplementary Figure 22e,f were calculated using OriginLab (2024b).

## Acknowledgement

This study was supported by the JC STEM Lab of Nanoscience and Nanomedicine funded by The Hong Kong Jockey Club Charities Trust, General Research Fund (RGC No. 17212424) and Early Career Scheme (RGC No. 27204623) from the Research Grants Council of Hong Kong SAR.

## Author contributions

F.W. conceived, designed and initiated the project. P.X. and F.W. developed the PL and AL imaging using TMOs and REOs. P.X. performed high-temperature N_2_/H_2_ reduction and characterizations of TMOs and REOs. S.X. contributed to the characterization of REOs. F.W.,

H.Y. and Z.L. designed and developed the scanning focused-ultrasound AL imaging system. F.W. designed and set up the spectral measurement system with ultrasound excitation. P.X. and Z.L. carried out the NIR-II PL imaging. P.X., Z.L., H.Y, S.X., Z.D., Z.W. and D.X. performed the *in vivo* NIR-II AL imaging. F.W. supervised the project and organized the collaboration. P.X. and F.W. wrote the manuscript. All authors contributed to the general discussion and revision of the manuscript.

## Competing interests

The authors declare no competing interests.

## Data availability

All data that support the findings of this study are presented in the main text and the Supplementary Information.

## Materials & Correspondence

Correspondence and requests for materials should be addressed to F.W.

## Notes

### Competing Interest Statement

The authors have declared no competing interest.

