## supplementary information for "Acoustoluminescence in Transition Metal and Rare Earth Oxides Beyond 1800 nm for In Vivo Imaging"

### SUPPLEMENTARY FIGURES

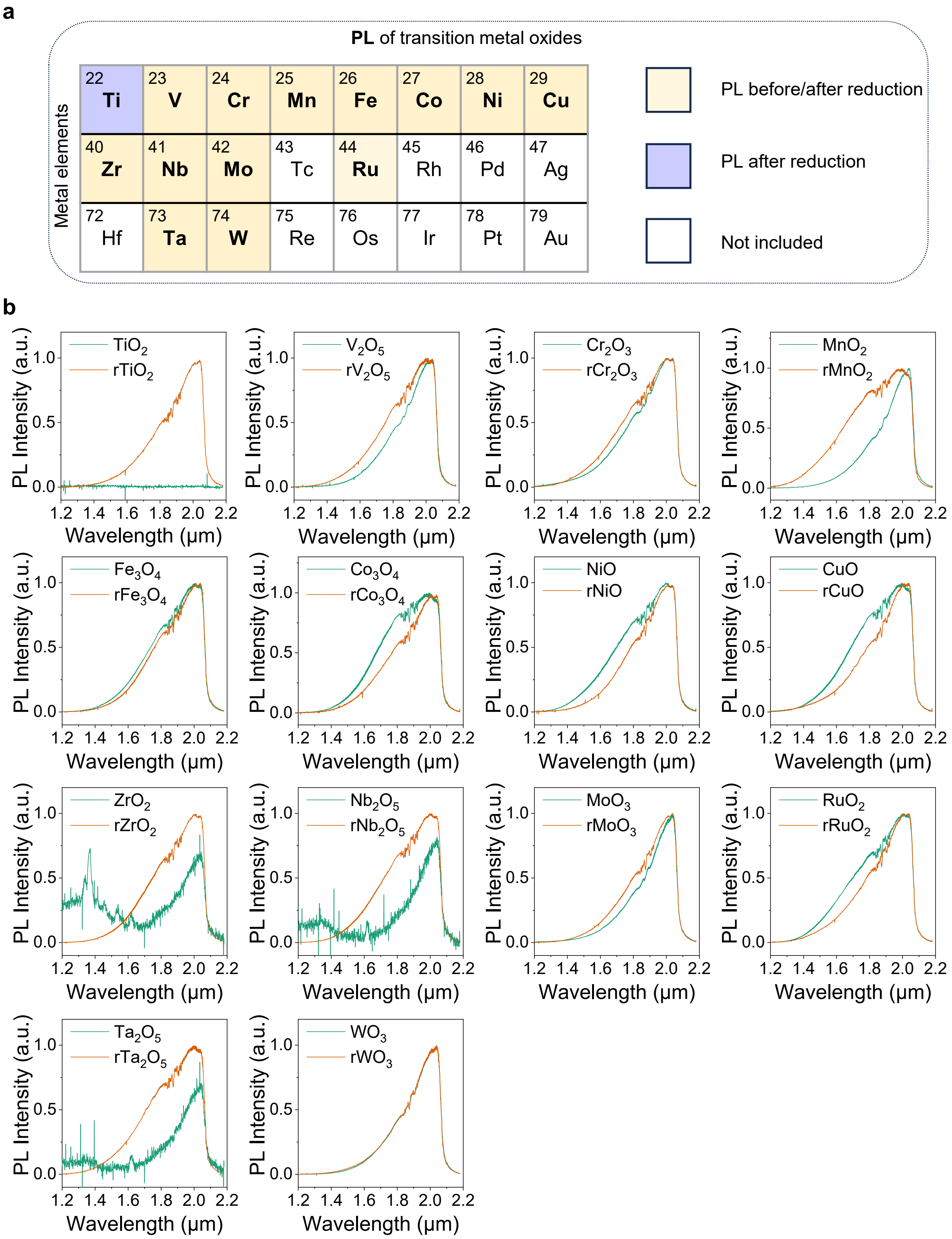


**Supplementary Figure 1. NIR-II PL in TMOs beyond 1800 nm.** (**a**) The effect of high-temperature $N_2/H_2$ reduction on NIR-II PL in TMOs. "PL before/after reduction" indicates that NIR-II PL emissions were observed both before and after $N_2/H_2$ reduction. "PL after reduction" means that NIR-II PL was only observed after the TMOs were subjected to $N_2/H_2$ reduction. These elements labeled in white were not investigated in this study. The reduction temperature was 1000 °C and the reduction time was 30 hours. To obtain powdered samples, $Co_3O_4$, NiO, and CuO were reduced at 600 °C. (**b**) NIR-II PL spectra of TMOs before and after $N_2/H_2$ reduction. NIR-II PL spectra were obtained using a focused 808-nm laser to excite powdered samples. The NIR-II spectrometer had an upper detection limit of ~2100 nm. The NIR-II PL intensities of $ZrO_2$, $Nb_2O_5$, and $Ta_2O_5$ can be enhanced by $N_2/H_2$ reduction.

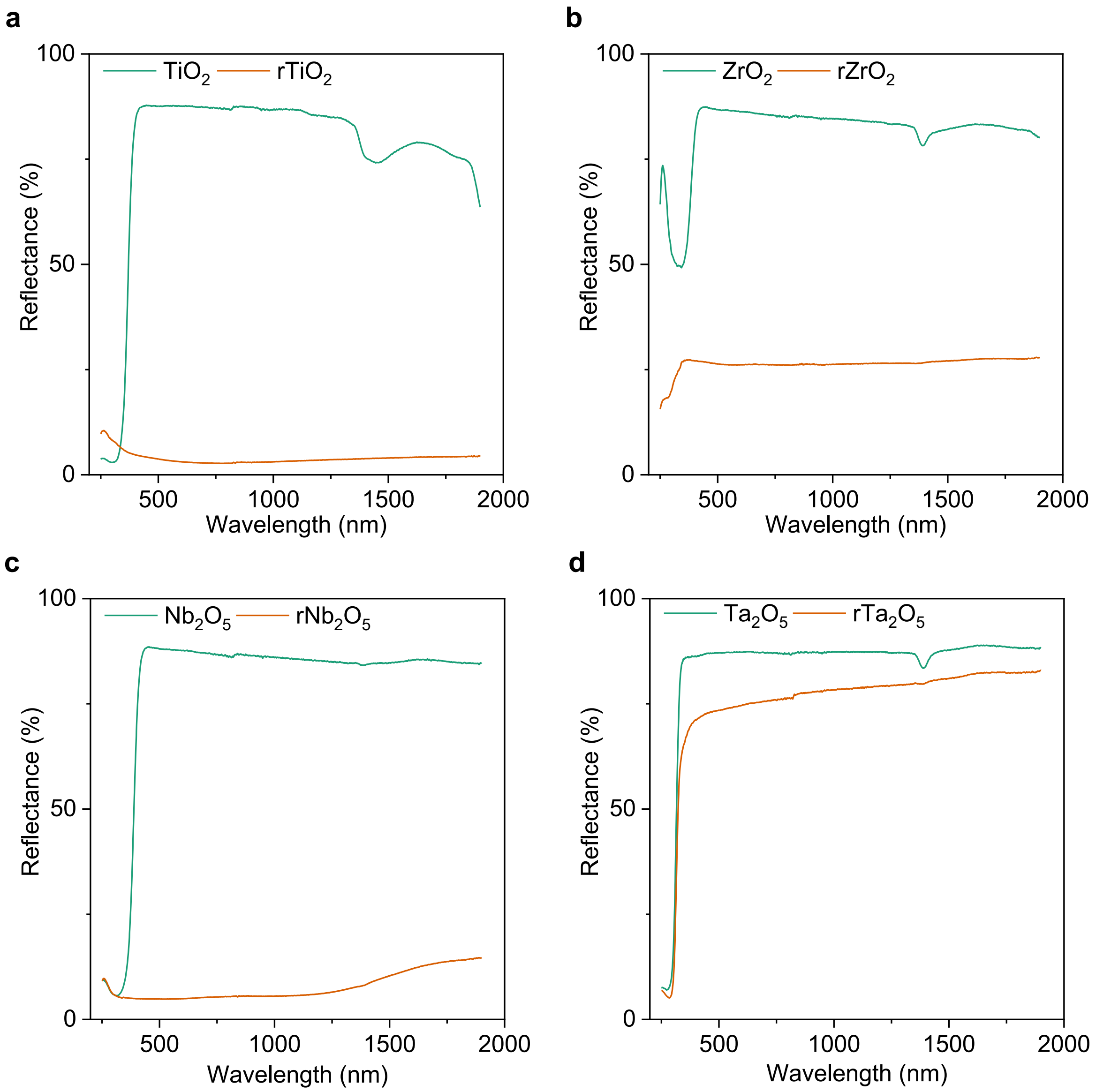


**Supplementary Figure 2. Diffuse reflectance spectra of $TiO_2$, $ZrO_2$, $Nb_2O_5$, and $Ta_2O_5$ before and after $N_2/H_2$ reduction.** The reduction was performed in an $N_2/H_2$ mixed gas at 1000 °C for 30 hours. The four TMOs without $N_2/H_2$ reduction had lower absorption at 808 nm, which may account for their weaker NIR-II PL intensities under a focused 808-nm laser excitation, compared to those of pristine samples, as shown in Supplementary Fig. 1.

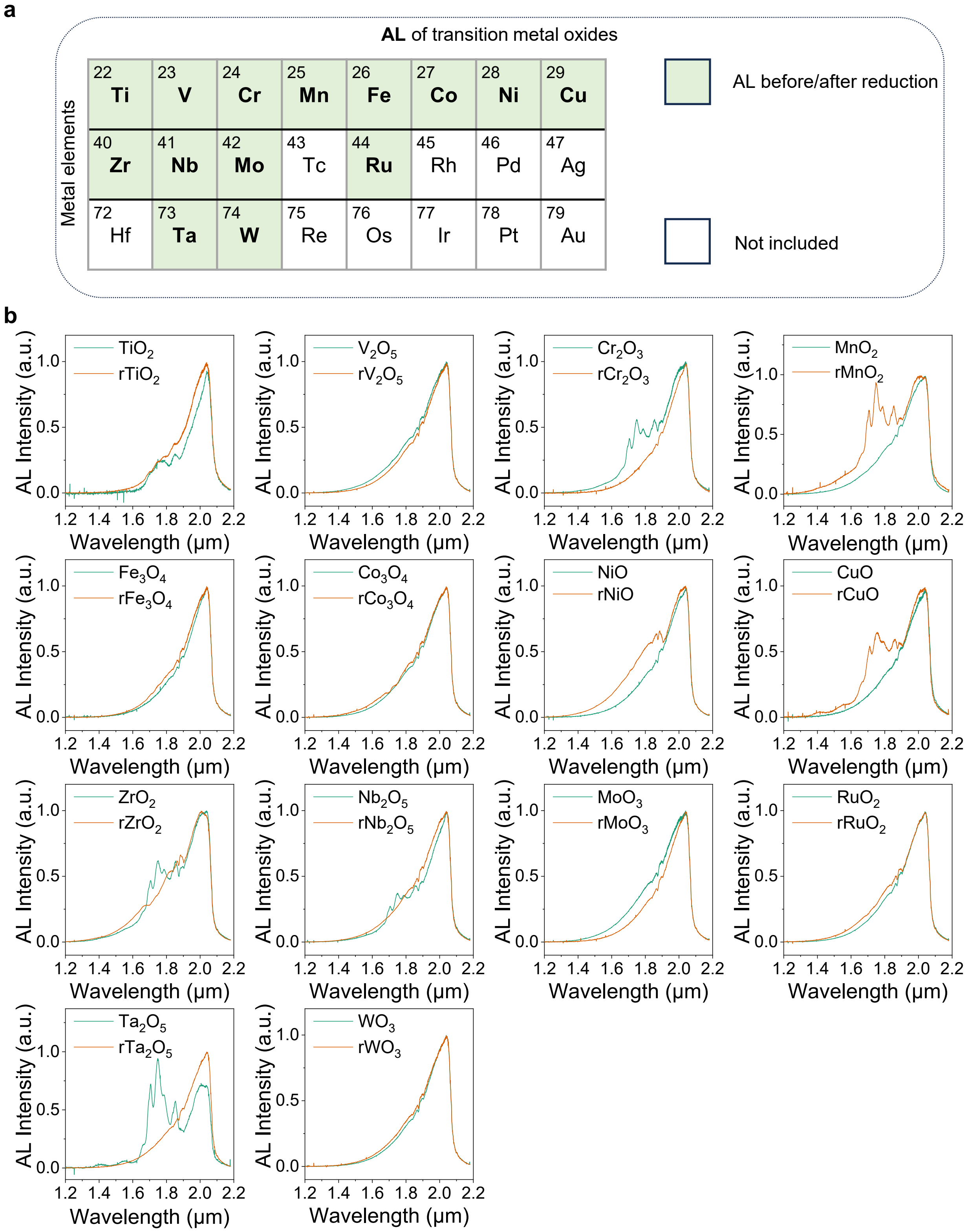


**Supplementary Figure 3. NIR-II AL in TMOs beyond 1800 nm.** (**a**) The influence of high-temperature $N_2/H_2$ reduction on NIR-II AL in TMOs. “AL before/after reduction” means that AL was observed in the TMOs both before and after $N_2/H_2$ reduction. The reduced TMOs are marked with “r”. $Co_3O_4$, NiO, and CuO were reduced at 600 °C for 30 hours, while other TMOs were reduced at 1000 °C for 30 hours. All pristine and reduced TMOs showed NIR-II AL emissions. White-colored elements were not included in this study. (**b**) AL spectra of TMOs before and after

$N_2/H_2$ reduction. The upper detection limit of the spectrometer was ~2100 nm. To record the NIR-II AL spectra, a ~0.5-mm-thick layer of powdered pristine or reduced TMOs was embedded at the bottom of PDMS substrates and excited by a 1 MHz ultrasonic therapy device. The AL spectra of $TiO_2$, $Cr_2O_3$, $MnO_2$, NiO, CuO, $ZrO_2$, $Nb_2O_5$, and $Ta_2O_5$ can be modulated by $N_2/H_2$ reduction.

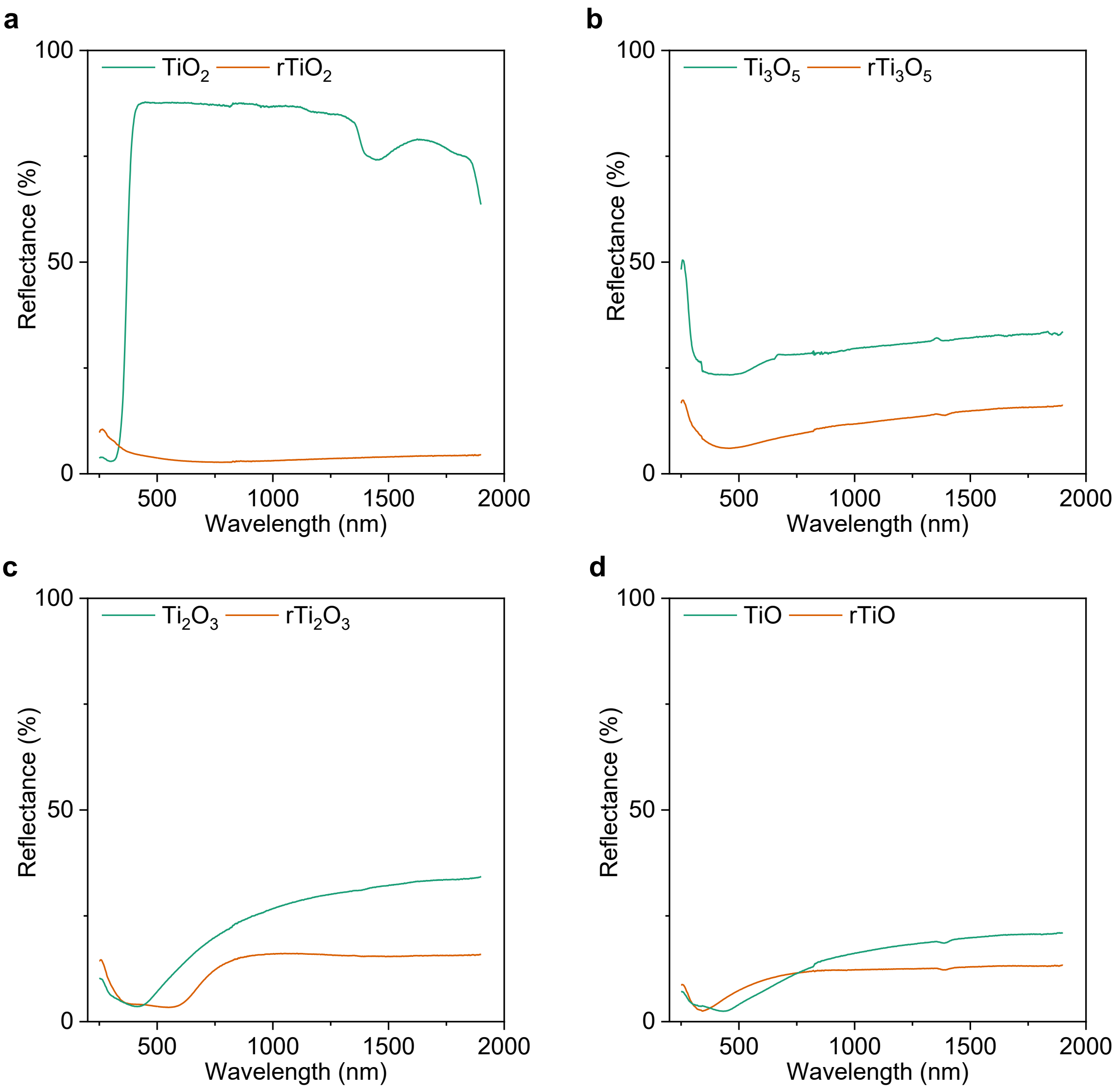


**Supplementary Figure 4. Diffuse reflectance spectra of $TiO_2$, $Ti_3O_5$, $Ti_2O_3$, and TiO before and after $N_2/H_2$ reduction.** The reduction temperature was 1000 °C, and the reduction time was 30 hours. Except for $TiO_2$ before reduction, the other titanium oxides had higher absorption at 808 nm, which may account for their obvious NIR-II PL intensities under a focused 808-nm laser, as shown in Figure 1g-i. $rTiO_2$ has the strongest absorption in the wavelength range of ~400-1900 nm, which explains its relatively higher NIR-II PL intensities among all four titanium oxides excited by the focused 808-nm laser.

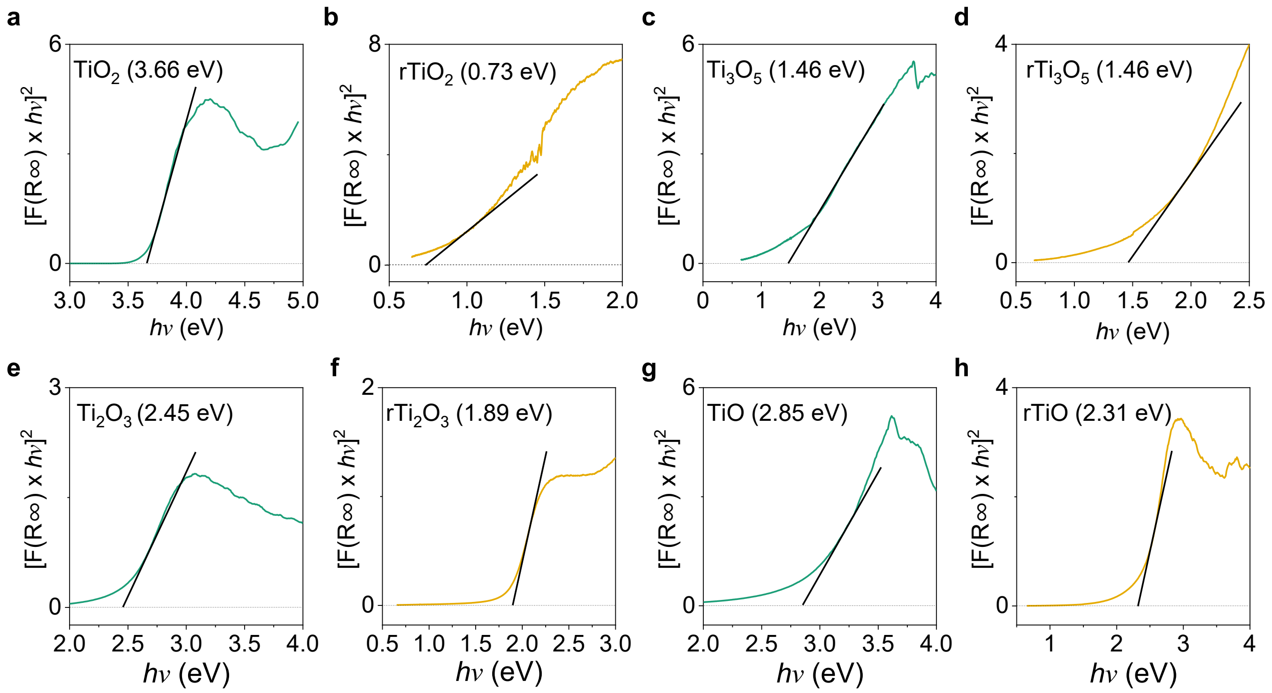


**Supplementary Figure 5. Bandgap difference of $TiO_2$, $Ti_3O_5$, $Ti_2O_3$, and TiO before and after $N_2/H_2$ reduction.** Plots of $[F(R\infty) \times h\nu]^2$ vs. $h\nu$ (eV) of (**a**) $TiO_2$, (**b**) $rTiO_2$, (**c**) $Ti_3O_5$, (**d**) $rTi_3O_5$, (**e**) $Ti_2O_3$, (**f**) $rTi_2O_3$, (**g**) TiO, and (**h**) rTiO, respectively. $rTiO_2$ (0.73 eV), reduced for 30 hours in $N_2/H_2$-mixed gas, exhibits the lowest bandgap value among titanium oxides. $TiO_2$ (3.66 eV) has the largest bandgap among the titanium oxides.

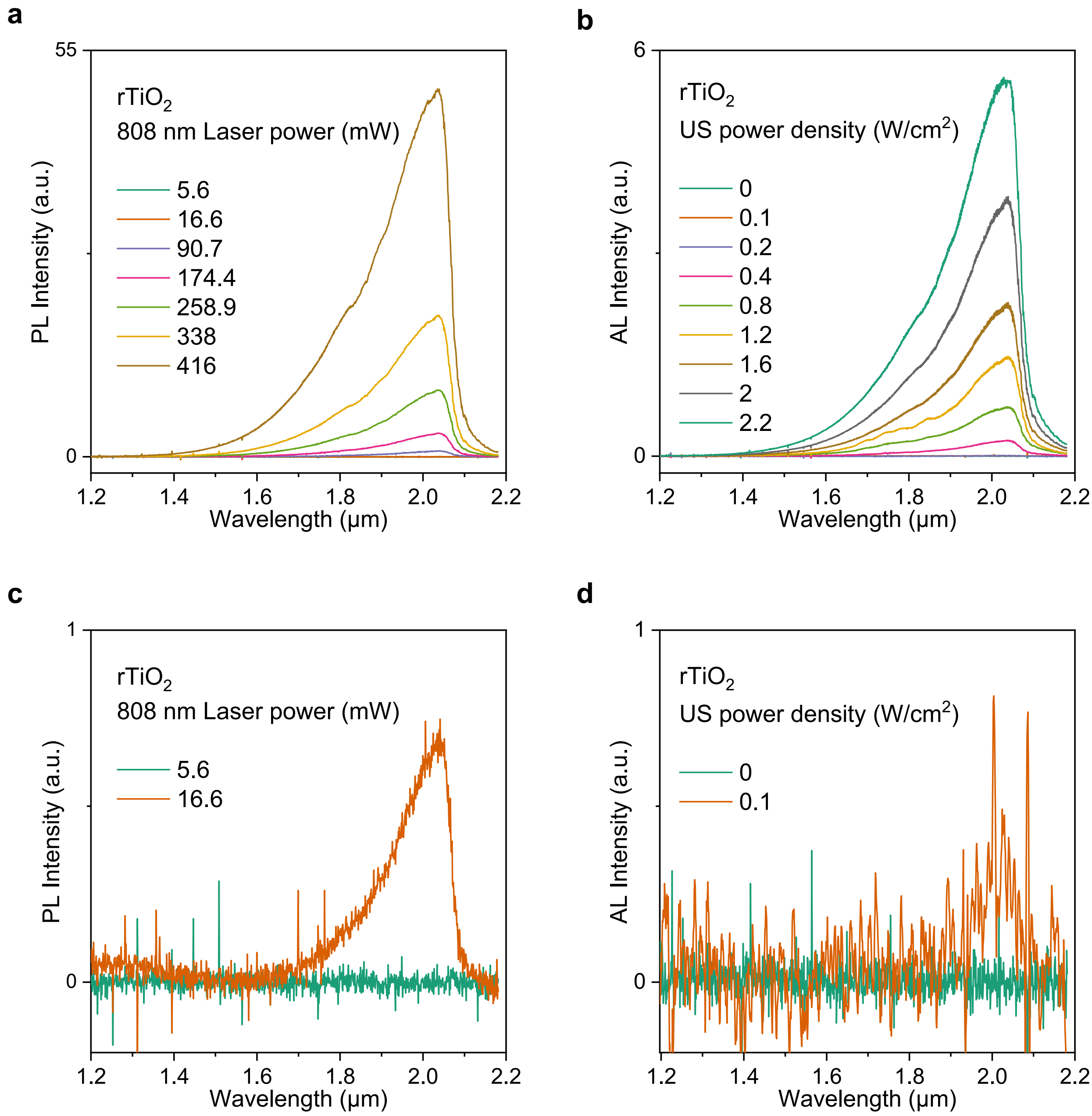


**Supplementary Figure 6. NIR-II PL and AL in $rTiO_2$ under different laser powers and US power densities.** (**a**) PL spectra of $rTiO_2$ excited by a focused 808-nm laser with different powers in the range of 5.6-416 mW. (**b**) AL spectra of $rTiO_2$ excited by a 1 MHz ultrasonic therapy device with different US power densities in the range of 0-2.2 $W/cm^2$. (**c**) PL spectra of $rTiO_2$ under focused 808-nm laser excitation at powers of 5.6 mW and 16.6 mW showed that the laser excitation threshold can be as low as 16.6 mW. (**d**) AL spectra of $rTiO_2$ at US power densities of 0 $W/cm^2$ and 0.1 $W/cm^2$ indicated that the ultrasound excitation threshold was ~0.1 $W/cm^2$. AL spectra at 0.1 and 0.2 $W/cm^2$ ultrasound power intensities were recorded ~15 s after turning on the ultrasonic therapy device, while AL spectra at higher ultrasound power intensities (0.4 - 2.2 $W/cm^2$) were obtained after waiting 10 s following ultrasonic stimulation. $rTiO_2$ was reduced at 1000 °C for 30 hours. The upper detection limit of the spectrometer was ~2100 nm.

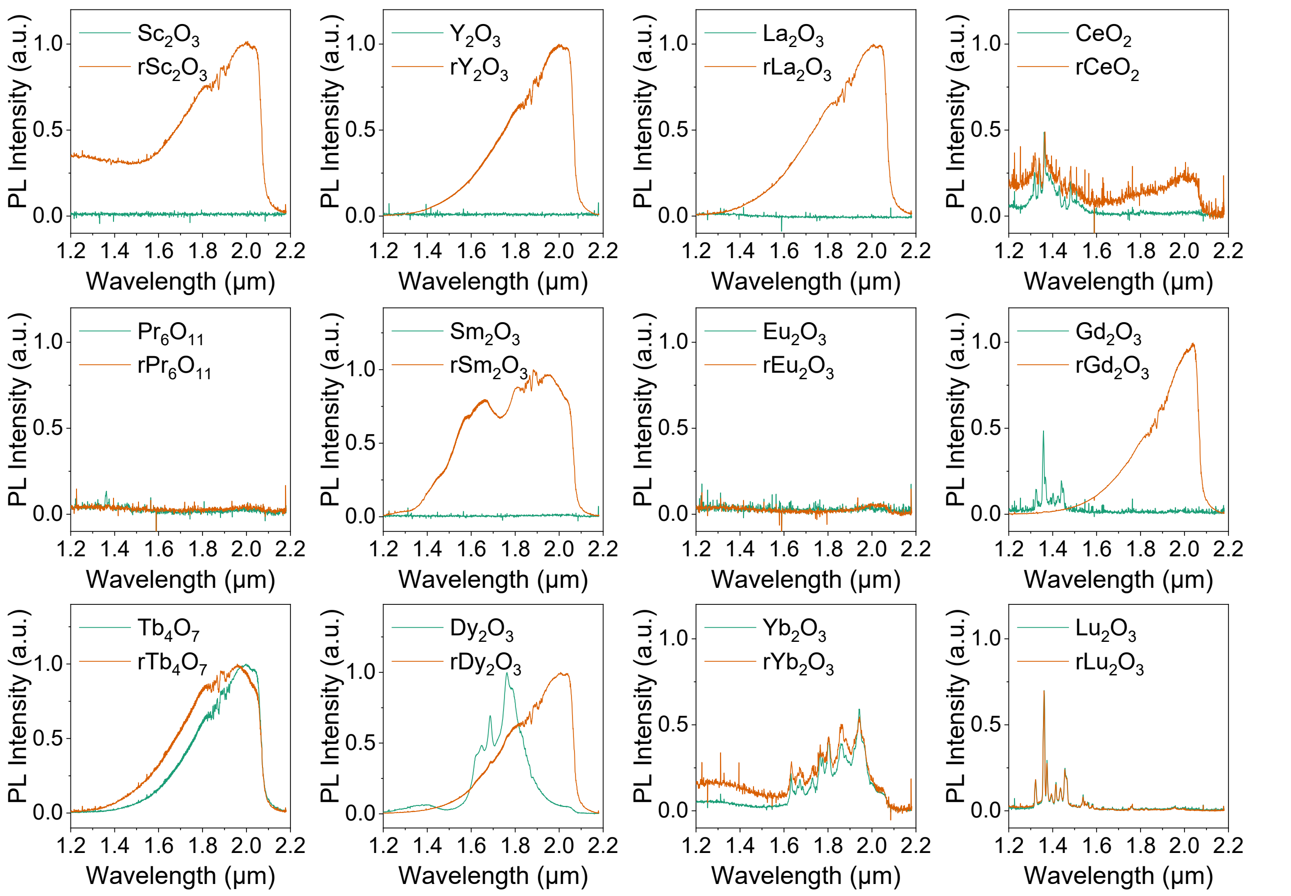


**Supplementary Figure 7. Non-conventional NIR-II PL spectra of REOs before and after $N_2/H_2$ reduction at 1000 °C for 30 hours**. The non-conventional NIR-II PL spectra of pristine and reduced $Nd_2O_3$, $Er_2O_3$, $Tm_2O_3$, and $Ho_2O_3$ were shown in Figure 2d. Firstly, $N_2/H_2$ reduction can reshape the spectral profiles of $Tb_4O_7$ and $Dy_2O_3$. Secondly, non-conventional NIR-II PL emissions in $Sc_2O_3$, $Y_2O_3$, $La_2O_3$, $Sm_2O_3$, and $Gd_2O_3$ could only be observed after $N_2/H_2$ reduction. Thirdly, only weak non-conventional NIR-II PL signals were observed in $CeO_2$, $Yb_2O_3$, and $Lu_2O_3$. Finally, no non-conventional NIR-II PL emissions were observed in $Pr_6O_{11}$ and $Eu_2O_3$ before and after $N_2/H_2$ reduction. Except for $Tm_2O_3$ and $rTm_2O_3$, which were excited with a focused 975-nm laser, all the rest spectra were measured by exciting the powdered samples with a focused 808-nm laser. An 1100-nm long-pass filter was used to filter out the laser light. The upper detection limit of the spectrometer was ~2100 nm. Pm related oxides were not discussed in this study.

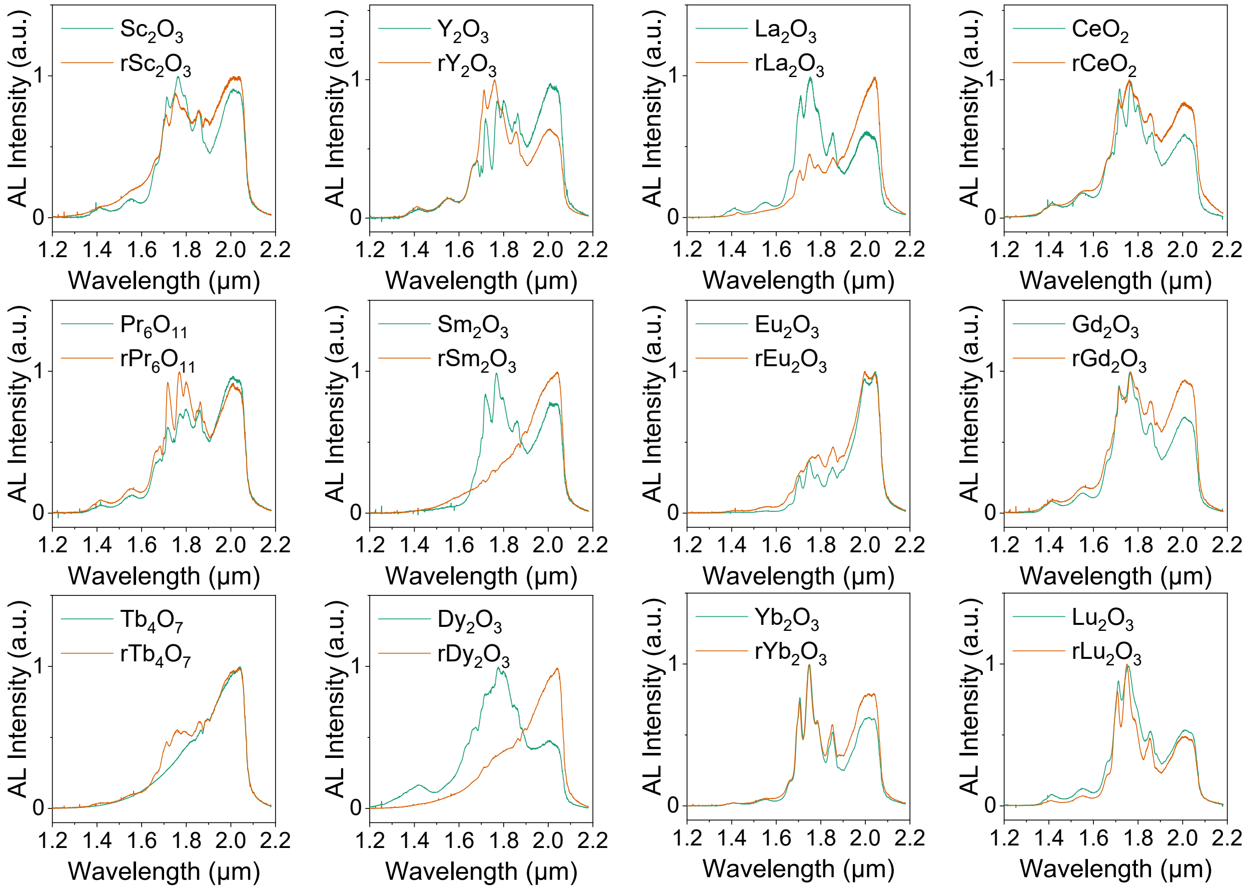


**Supplementary Figure 8. AL spectra of REOs before and after $N_2/H_2$ reduction at 1000 °C for 30 hours**. The NIR-II AL spectra of pristine and reduced $Nd_2O_3$, $Er_2O_3$, $Tm_2O_3$, and $Ho_2O_3$ were shown in Figure 2e. To obtain the NIR-II AL spectra of REOs, a ~0.5-mm layer of pristine or reduced REOs was placed at the bottom of a PDMS substrate and then excited by an ultrasonic therapy device operating at 1 MHz. The upper detection limit of the spectrometer was ~2100 nm. $N_2/H_2$ reduction reshaped the NIR-II AL spectra of REOs after reduction. Obvious changes in NIR-II AL profiles were observed in $La_2O_3$, $Sm_2O_3$, and $Dy_2O_3$.

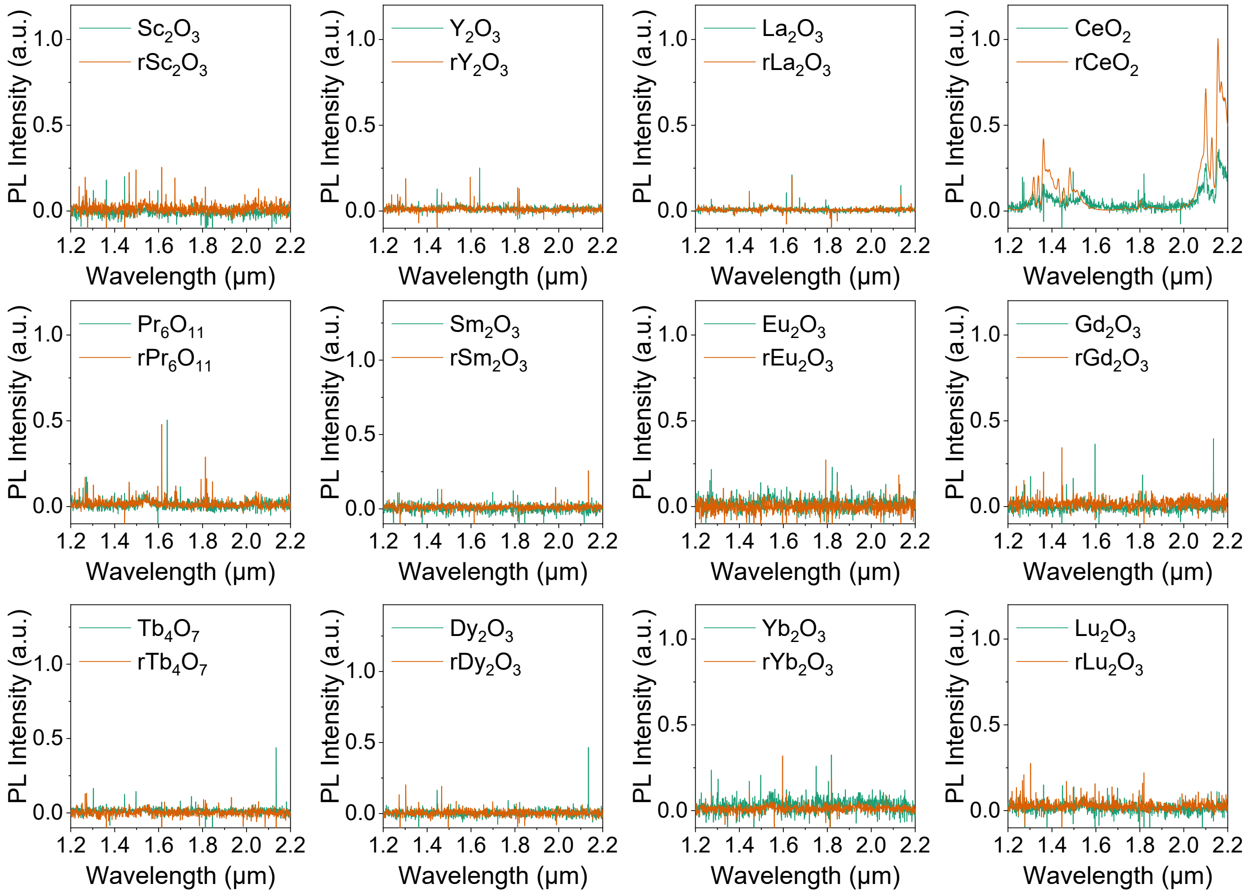


**Supplementary Figure 9. Intrinsic PL spectra of REOs before and after $N_2/H_2$ reduction at 1000 °C for 30 hours**. The obvious intrinsic NIR-II PL spectra of pristine and reduced $Nd_2O_3$ ($Nd^{3+}$, ~1370 nm, $^4F_{3/2}\rightarrow{}^4I_{13/2}$; ~1430 nm, $^4F_{3/2}\rightarrow{}^4I_{13/2}$), $Er_2O_3$ ($Er^{3+}$, ~1550 nm, $^4I_{13/2}\rightarrow{}^4I_{15/2}$), $Tm_2O_3$ ($Tm^{3+}$, ~1943 nm, $^3F_4\rightarrow{}^3H_6$), and $Ho_2O_3$ ($Ho^{3+}$, ~1213 nm, $^5I_6\rightarrow{}^5I_8$; ~2020 nm, $^5I_7\rightarrow{}^5I_8$) were shown in Figure 2c. A 365-nm UV LED was used to excite intrinsic PL from powdered samples. Obvious intrinsic NIR-II PL emissions in 1200-2200 nm were observed in $Nd_2O_3$, $Er_2O_3$, $Tm_2O_3$, $Ho_2O_3$, and $CeO_2$. Such intrinsic NIR-II PL emissions from pristine and reduced $CeO_2$ may be due to impurities in the raw materials or other unknown factors, and no similar results have been reported previously. The upper detection limit of the spectrometer was ~2100 nm.

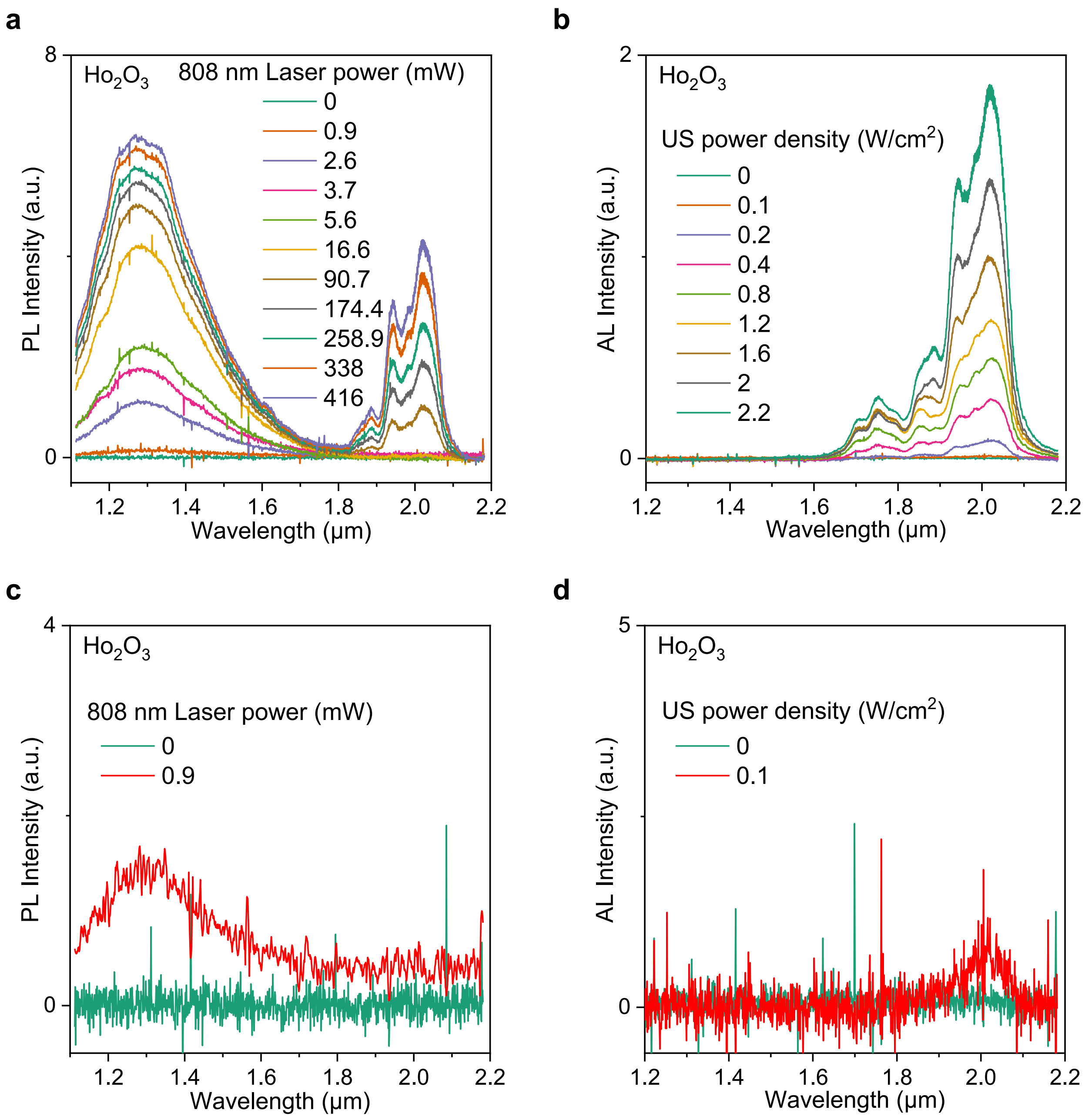


**Supplementary Figure 10. NIR-II PL and AL spectra of $Ho_2O_3$ under different laser powers and ultrasound (US) power densities.** (**a**) PL spectra of $Ho_2O_3$ under a focused 808-nm laser at different laser powers (0-416 mW). (**b**) AL spectra of $Ho_2O_3$ under excitation from 1 MHz ultrasonic therapy device at different US power densities (0-2.2 W/cm$^2$). (**c**) PL spectra of $Ho_2O_3$ under 808-nm laser excitation at powers of 0 mW and 0.9 mW showed that the laser excitation threshold could be as low as 0.9 mW. (**d**) AL spectra of $Ho_2O_3$ under ultrasound excitation at power densities of 0 and 0.1 W/cm$^2$ indicated that the US excitation threshold was ~0.1 W/cm$^2$. The minimum power density for this ultrasonic therapy device was 0.1 W/cm$^2$. AL spectra at 0.1 and 0.2 W/cm$^2$ ultrasound power intensities were recorded ~15 s after turning on the ultrasonic therapy device, while other AL spectra (0.4-2.2 W/cm$^2$) were obtained after waiting for 10 s following ultrasonic stimulation. The upper detection limit of the spectrometer was ~2100 nm.

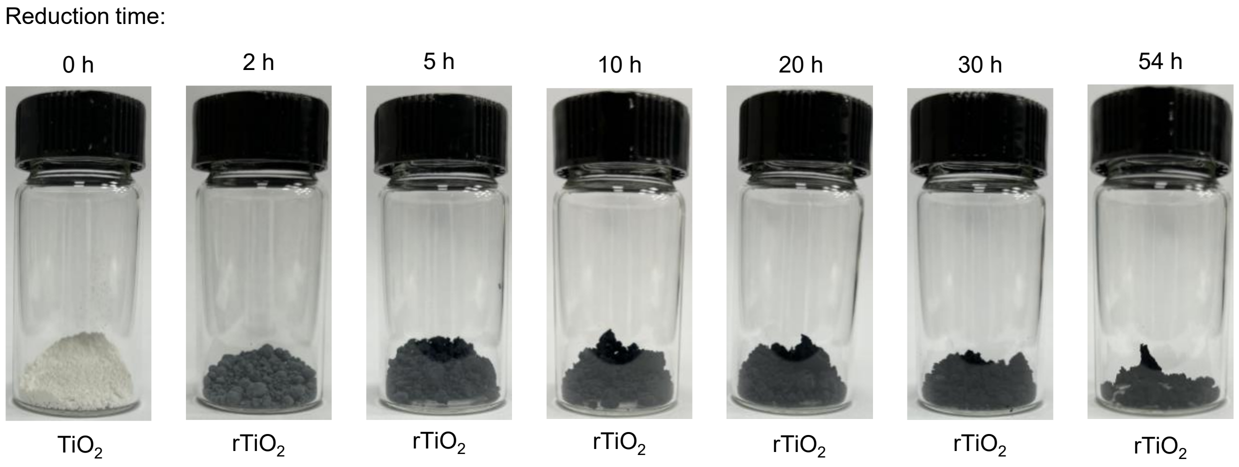


**Supplementary Figure 11. Photographs of $rTiO_2$ under different reduction times.** The reduction temperature was 1000 °C (see details in Methods). The color changed from white (0 hours), to light black (2 and 5 hours), and then to dark black (10, 20, 30, and 54 hours). Reduced $TiO_2$ appeared dark due to their high absorption in visible window.

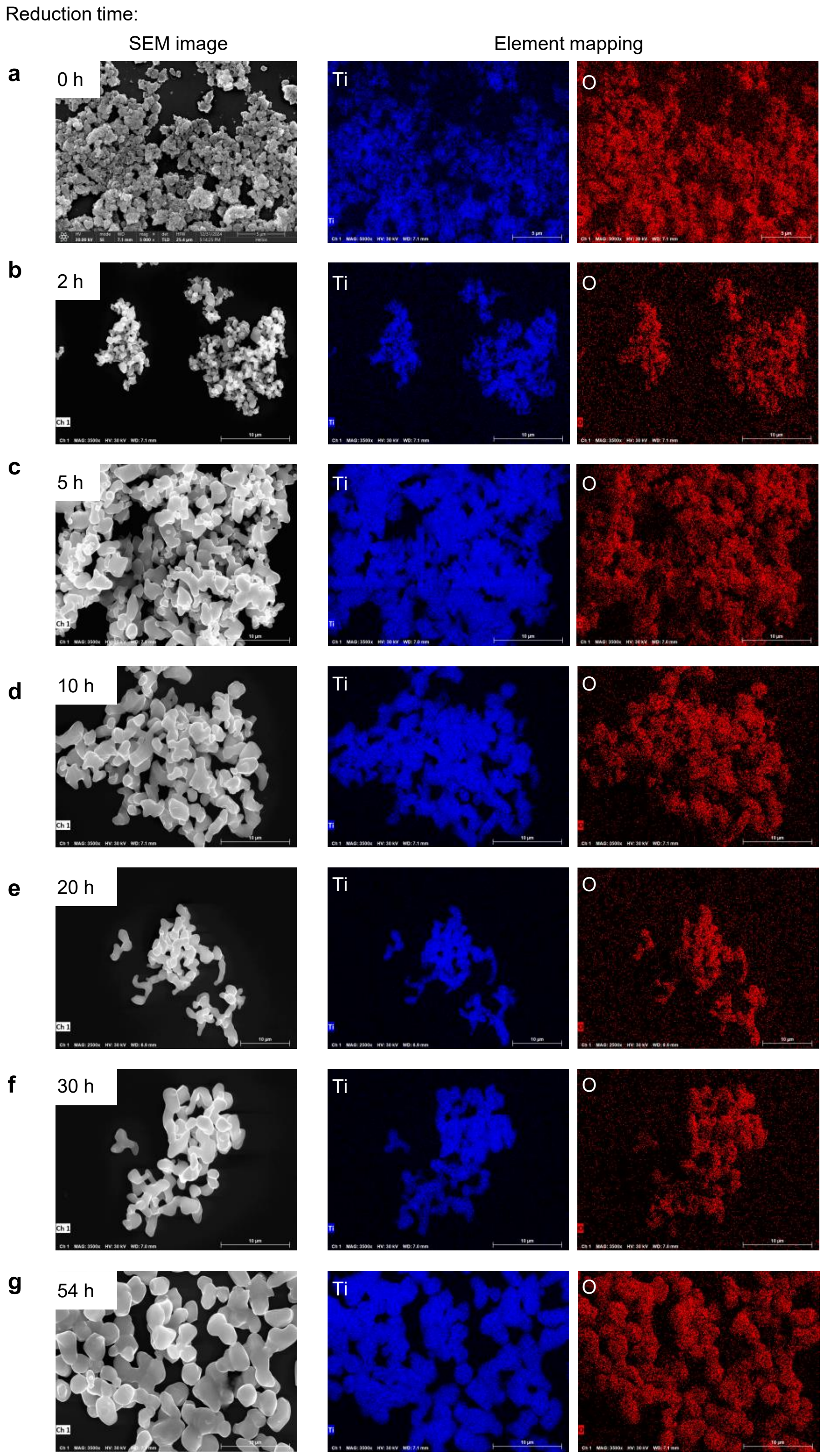


**Supplementary Figure 12. SEM images of $rTiO_2$ under different reduction times.** The reduction temperature was 1000 °C (see details in Methods). The size of raw $TiO_2$ was in the range of 5-10 nm, and its grain profile could not be resolved in (**a**). (**b**) The grain outline became visible in the 5-hour-reduced $TiO_2$. (**c-g**) Further extending the reduction time resulted in larger $rTiO_2$ with average grain sizes of several micrometers.

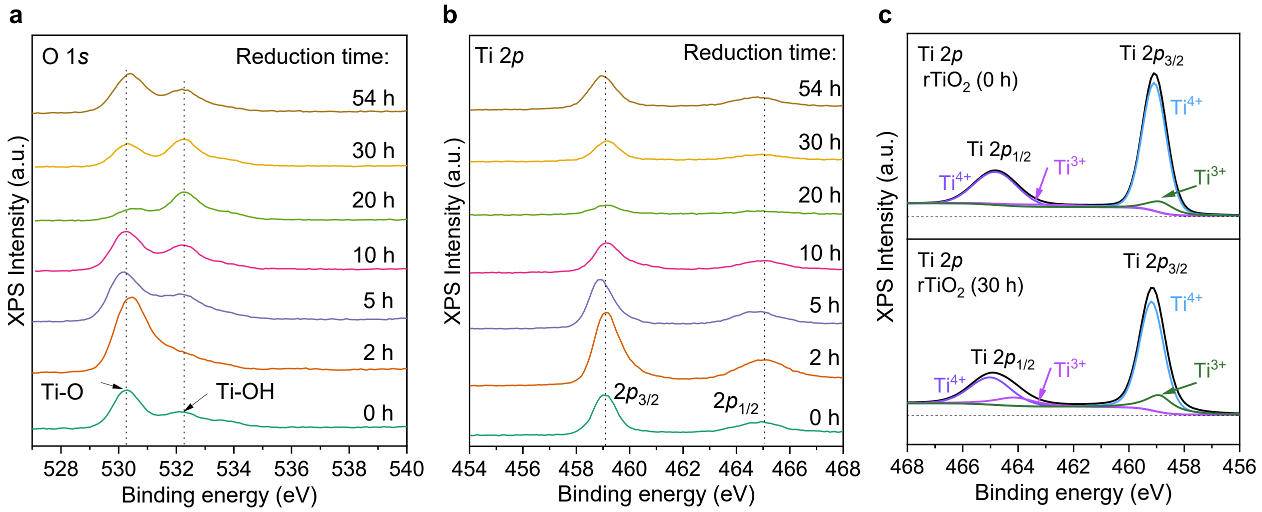


**Supplementary Figure 13. O 1*s* and Ti 2*p* XPS spectra of r$TiO_2$ under different reduction times**. (**a**) O 1*s* XPS spectra of $TiO_2$ under different reduction times (0-54 hours). All samples were reduced at 1000 °C in a $N_2/H_2$ mixed gas. As the reduction time increased, the peak intensity of Ti-OH gradually approached that of Ti-O, becoming equal after 30 hours of reduction, and dropped below that of Ti-O after 54 hours. (**b**) Two Ti 2*p* XPS peaks at ~459.1 eV and 465.1 eV were observed, which corresponded to $2p_{3/2}$ and $2p_{1/2}$ of Ti, respectively. (**c**) Ti 2*p* XPS spectra of $TiO_2$ before and after 30 hours of reduction. Low-valence titanium ($Ti^{3+}$) can be observed even before reduction, which is due to the presence of intrinsic $V_o$.[1] High-temperature $N_2/H_2$ reduction can induce more $V_o$ (Figure 3f), along with the formation of additional $Ti^{3+}$ ions.

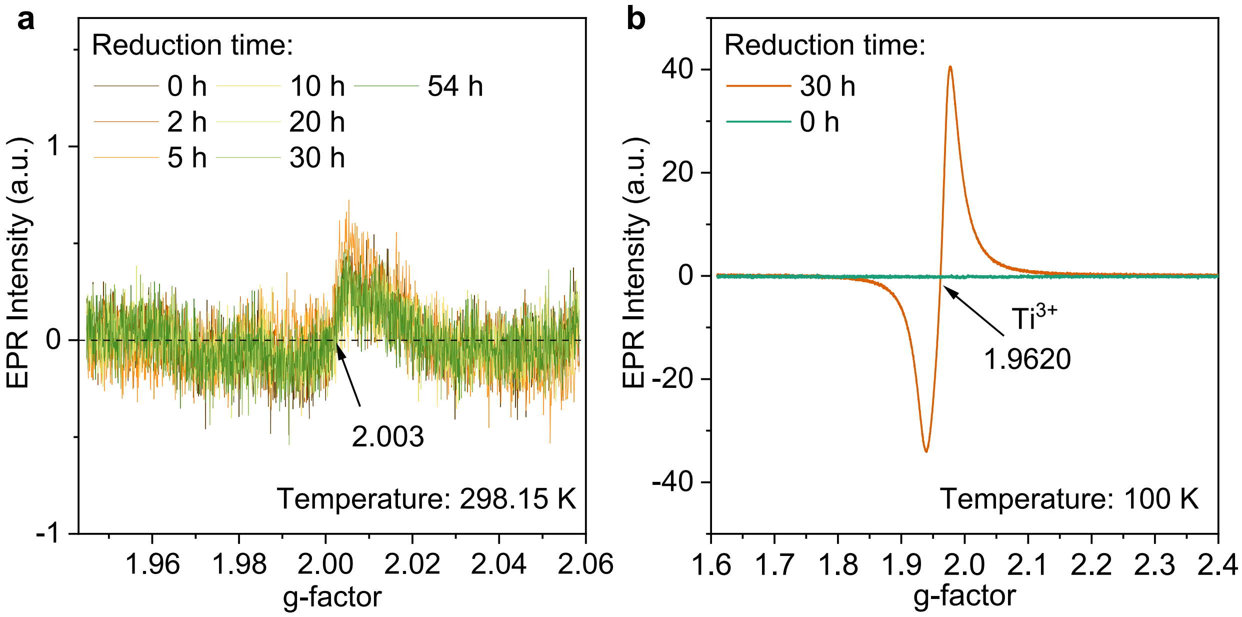


**Supplementary Figure 14.** (**a**) Room-temperature (298.15 K) EPR spectra of $rTiO_2$ after different reduction times of 0, 2, 5, 10, 20, 30 and 54 hours, respectively. $N_2/H_2$-mixed gas was used to reduce powdered samples at 1000 °C. The hydroxyl group was inferred to be formed on the surface[2], while $V_o$ may be present in the subsurface, which could not be easily detected due to the limit detection depth of EPR and the potential influence of surface hydroxyl groups, as indicated by faint EPR signals at room temperature. (**b**) Low temperature (100 K) EPR spectra of $rTiO_2$ reduced for 0 hours and 30 hours. We observed an obvious $Ti^{3+}$ EPR signal (g-factor = 1.9620) in $rTiO_2$ reduced for 30 hours[3, 4], while this signal was absent in pristine $TiO_2$. The $V_o$ (g-factor = 2.003) signal was completely obscured by the strong $Ti^{3+}$ signal. The low-temperature EPR signal confirmed the existence of $Ti^{3+}$-$V_o$ associates[4].

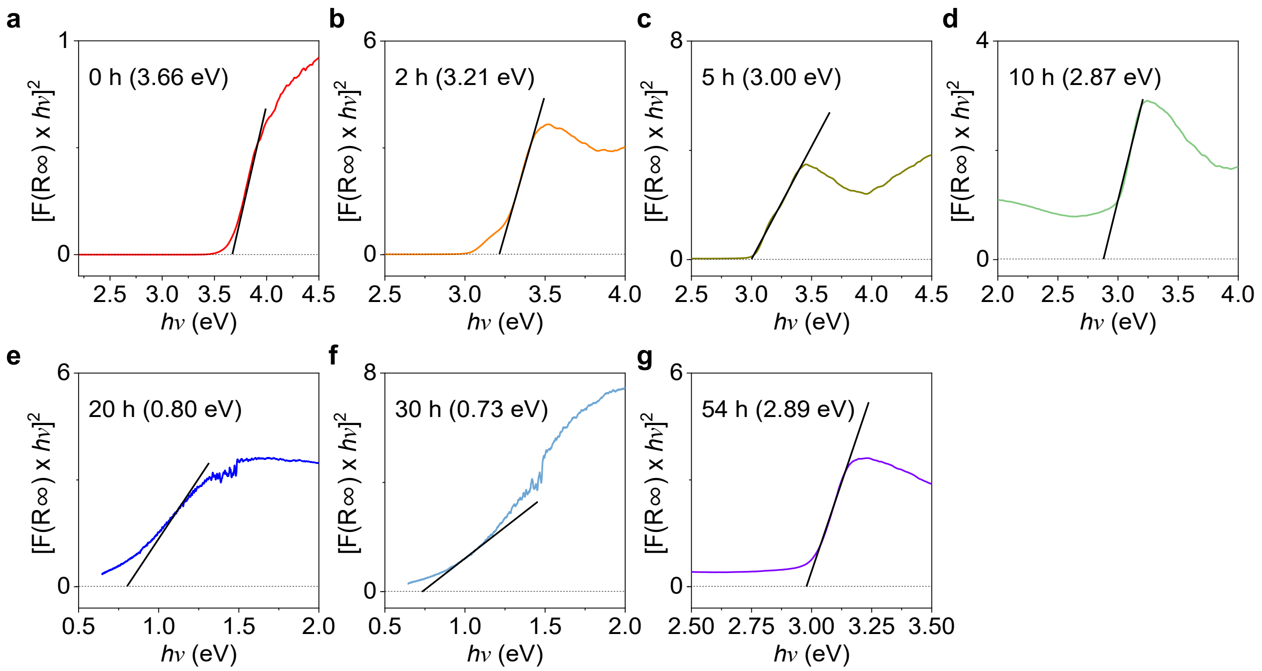


**Supplementary Figure 15. Bandgap of $TiO_2$ before and after reduction in $N_2/H_2$-mixed gas at 1000 °C for different reduction times.** Plot of $[F(R\infty) \times h\nu]^2$ vs. $h\nu$ of $TiO_2$ reduced for (**a**) 0 hours, (**b**) 2 hours, (**c**) 5 hours, (**d**) 10 hours, (**e**) 20 hours, (**f**) 30 hours, and (**g**) 54 hours. As the reduction time increased from 0 to 30 hours, the bandgap value decreased, reaching a minimum of 0.73 eV at 30 hours. A narrower bandgap requires less energy to excite electrons from the valence band to the conduction band. With the introduction of more intermediate gap states, more carriers may escape from defects and participate in the AL or PL emission processes[5].

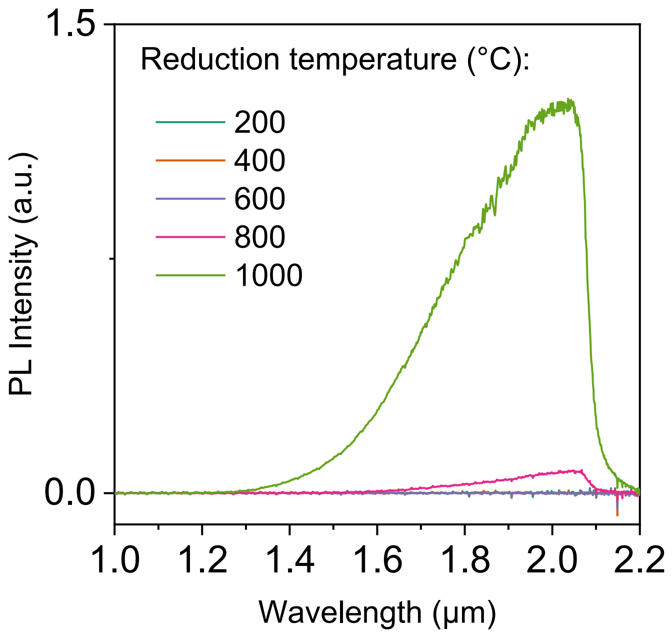


**Supplementary Figure 16. Influence of the reduction temperature on the NIR-II PL intensities of r$TiO_2$**. The reduction was performed in a flowing $N_2$/$H_2$-mixed gas for 30 hours at different temperatures ranging from 200 to 1000 °C. To obtain the NIR-II PL spectra, a focused 660-nm laser was used to excite powdered samples. An 1100-nm long-pass filter was used, and the upper detection limit of the spectrometer was ~2100 nm. Increasing the reduction temperature in the range of 200-1000 °C enhanced its NIR-II PL intensity.

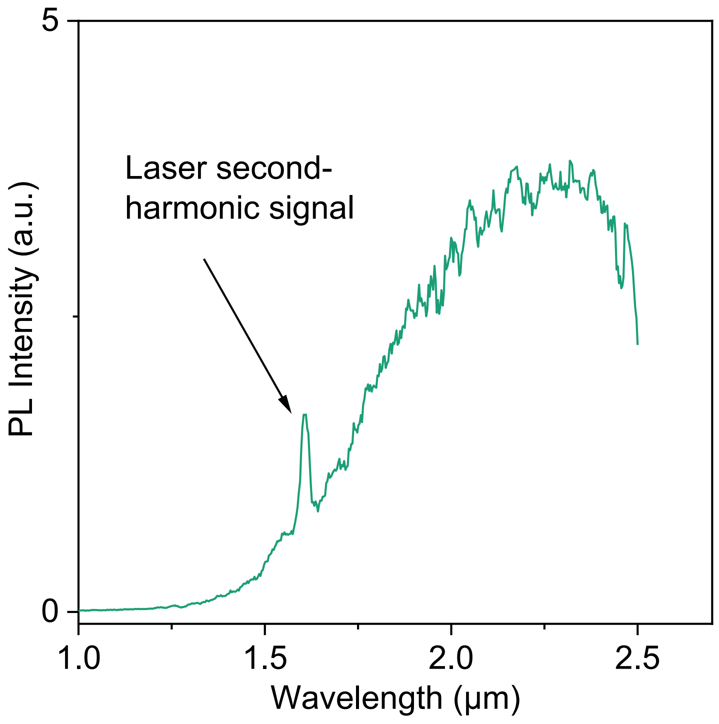


**Supplementary Figure 17.** The NIR-II PL spectrum of $TiO_2$ reduced for 30 hours was measured using an InGaAs-based spectrometer (NIR QUEST+2.5, Ocean Optics) with an upper detection limit of ~2500 nm. A broadband NIR-II PL spectrum peaked at ~2280 nm was observed. A focused 808-nm laser was used for excitation.

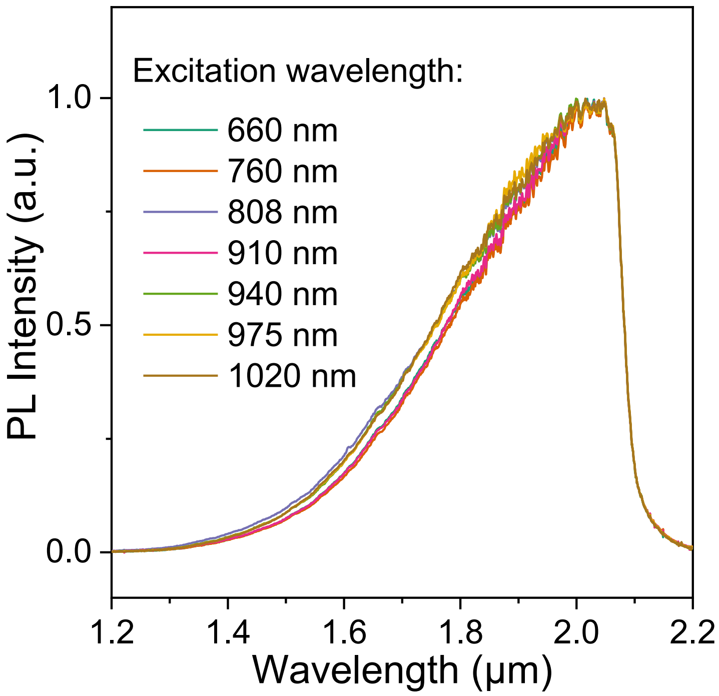


**Supplementary Figure 18.** NIR-II PL spectra of r$TiO_2$ powders under focused lasers with different excitation wavelengths of 660, 760, 808, 910, 940, 975, and 1020 nm, respectively. The powdered sample was obtained by reducing $TiO_2$ in $N_2/H_2$-mixed gas at 1000 °C for 30 hours. An 1100-nm long-pass filter was used. The upper detection limit of the spectrometer was ~2100 nm.

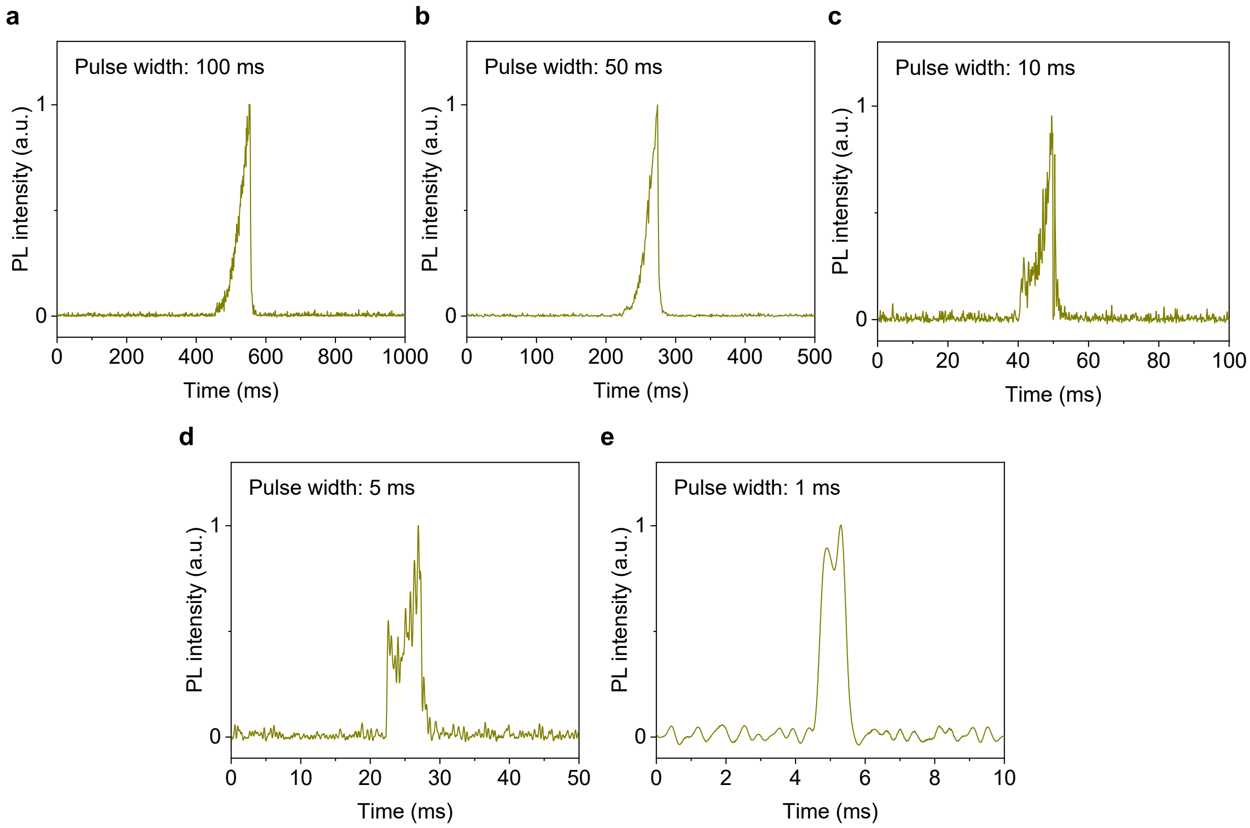


**Supplementary Figure 19.** Transient response of PL emission from $rTiO_2$ reduced for 30 hours under pulsed laser with a duration in the range of 1-100 ms. The PL was collected by an InGaAs PMT after passing through a 1500-nm long-pass filter.

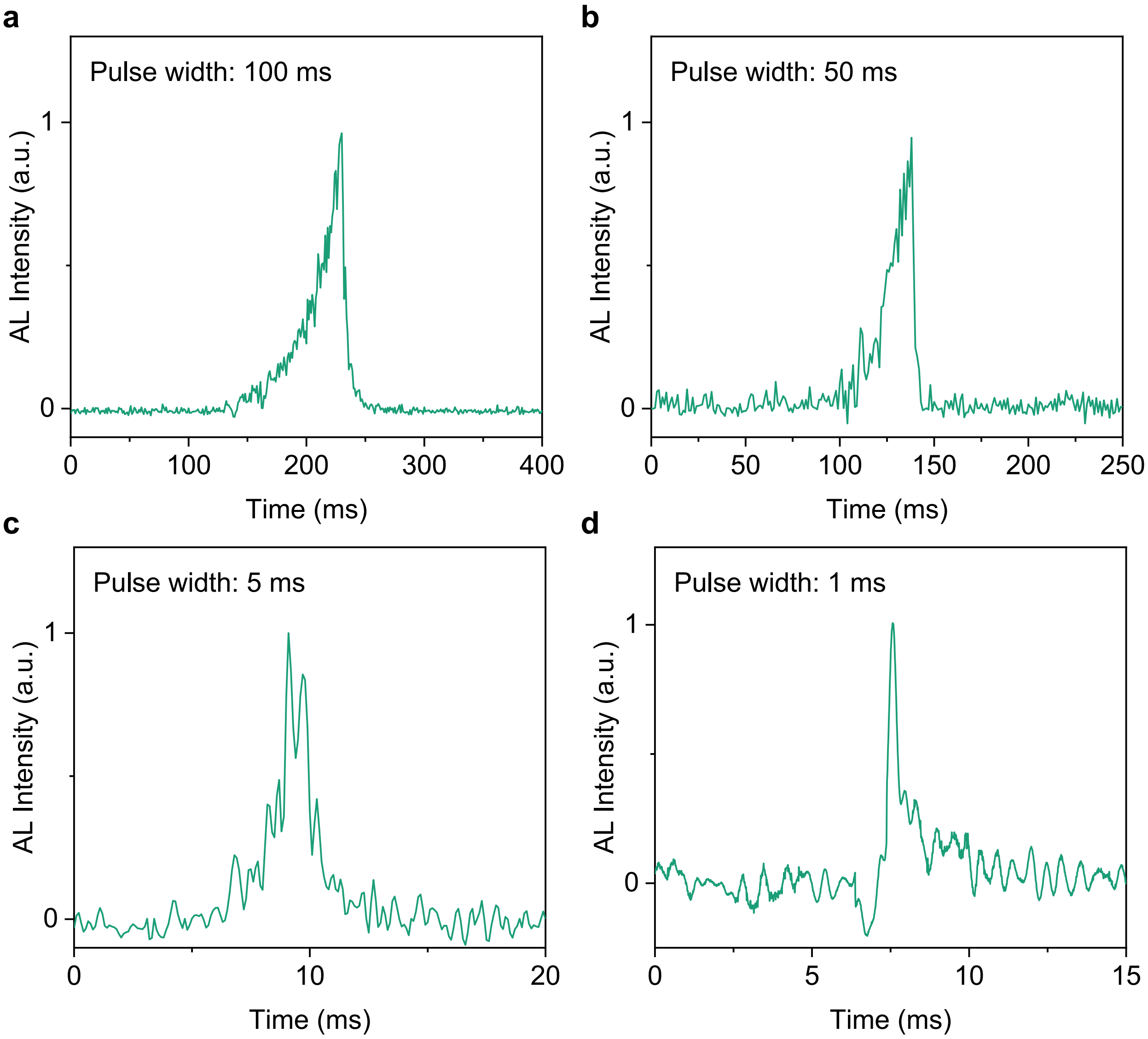


**Supplementary Figure 20. Transient response of AL emission from r$TiO_2$ reduced for 30 hours under pulsed ultrasound with a duration of 1-100 ms.** r$TiO_2$ was prepared by $N_2/H_2$ reduction of $TiO_2$ at 1000 °C for 30 hours. A ~0.5-mm-thick layer of r$TiO_2$ was placed at the bottom of a PDMS substrate and then excited by a focused ultrasound transducer operating at ~4.55 MHz. The AL was collected by an InGaAs PMT after passing through a 1500-nm long-pass filter.

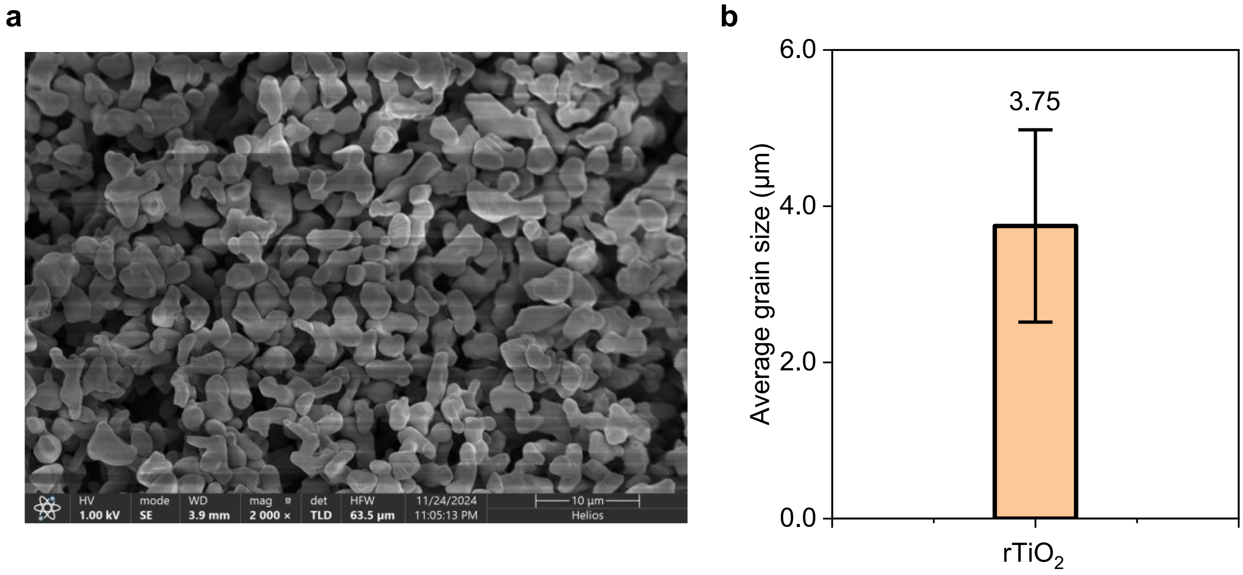


**Supplementary Figure 21.** (**a**) SEM image and (**b**) average grain size of $rTiO_2$ reduced for 30 hours at 1000 °C. Data are shown as mean ± s.d. derived from analyzing 20 measured values.

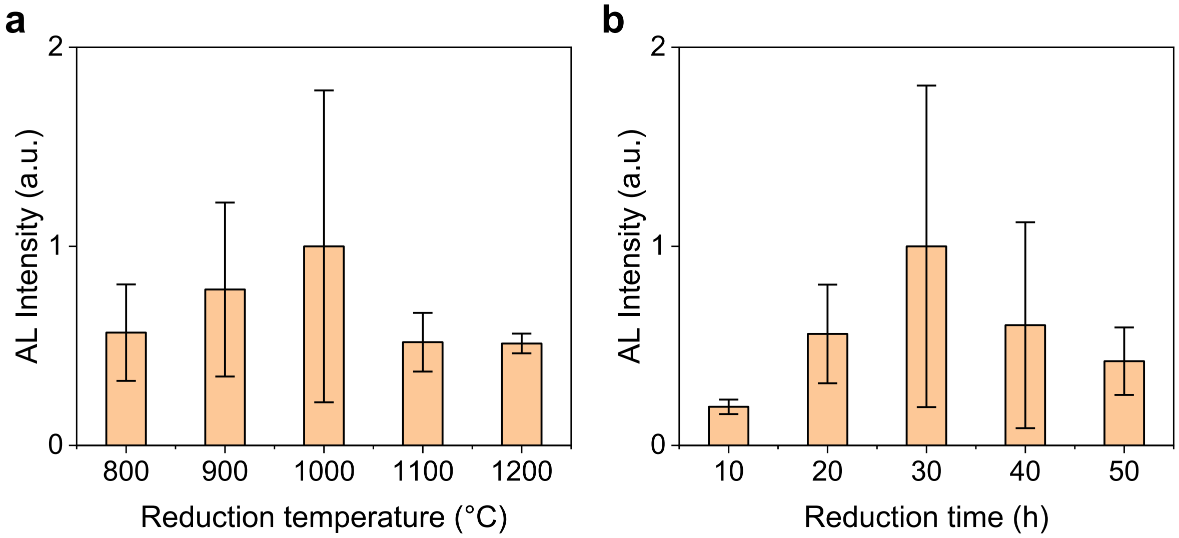


**Supplementary Figure 22. Optimization of NIR-II AL brightness in $rTiO_2$-MSN.** (**a**) NIR-II AL brightness of $rTiO_2$-MSN at different reduction temperatures (800, 900, 1000, 1100, and 1200 °C). The reduction time was 30 hours, and the MSN to $TiO_2$ mass ratio was 2:1. (**b**) Influence of reduction time on NIR-II AL brightness of $rTiO_2$-MSN. The reduction temperature was 1000 °C and the MSN to $TiO_2$ mass ratio was 2:1. To compare the NIR-II AL brightness, a ~0.5-mm-thick layer of powdered $rTiO_2$-MSN sample was embedded at the bottom of a PDMS substrate and positioned at the focused plane of an ultrasonic transducer operating at ~4.55 MHz. The material surface was positioned at the bottom, and NIR-II AL signals were recorded by an NIR-II camera after passing through the PDMS substrate. In (**a**) and (**b**), data were shown as mean ± s.d. derived from analyzing ~436 and 249 target data, respectively.

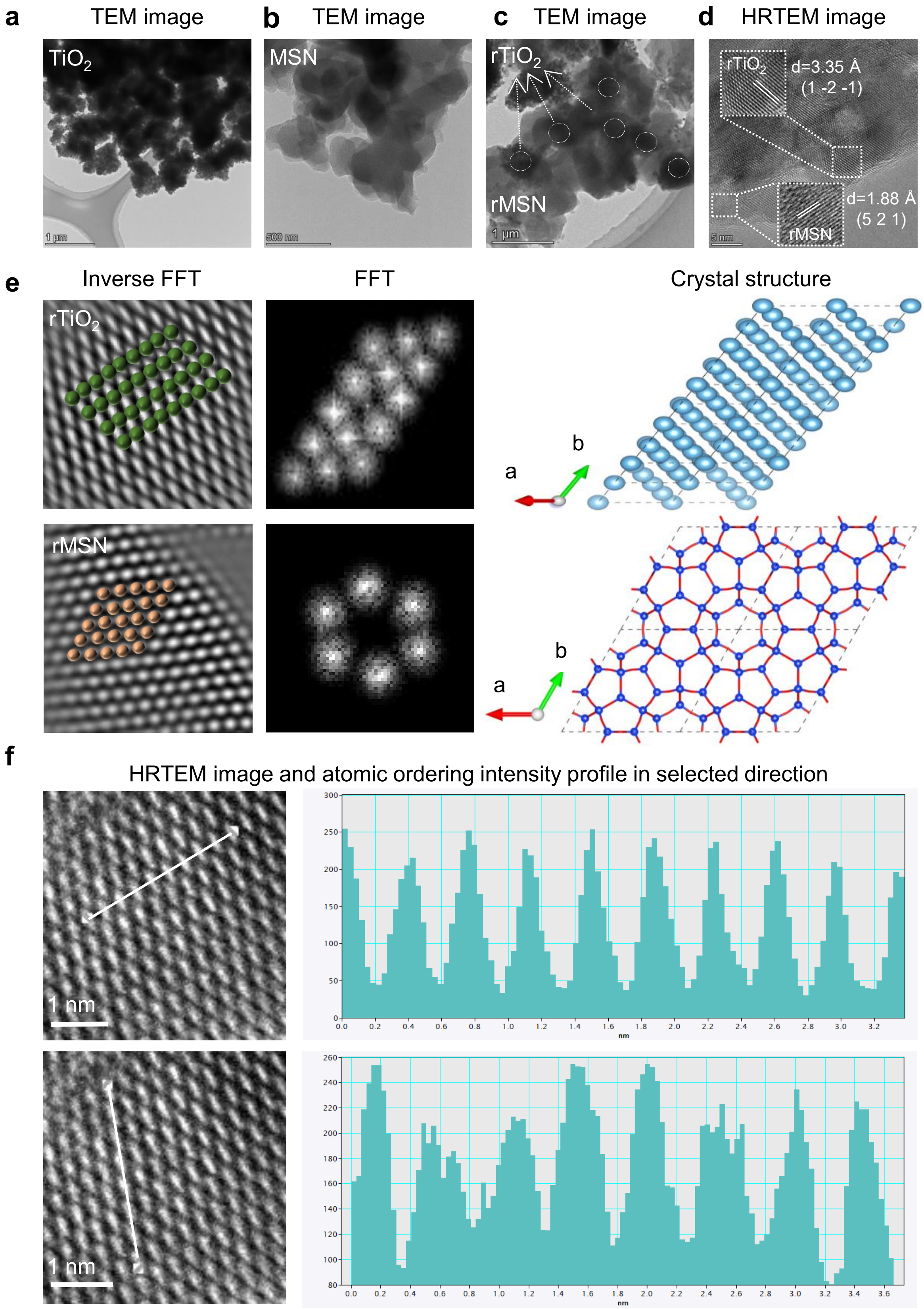


**Supplementary Figure 23. TEM characterization of $rTiO_2$-MSN.** (**a**) TEM image of $TiO_2$ raw material (0 hours, anatase, 01-089-4921). (**b**) TEM image of MSN raw material (P6/mmm, ICSD#48153). (**c**) TEM image of $rTiO_2$-MSN reduced by $N_2/H_2$-mixed gas at 1000 °C for 30 hours, with an MSN to $TiO_2$ ratio of 2:1. (**d**) HRTEM image of $rTiO_2$-MSN shown in (**c**). The interplanar spacing of $rTiO_2$ was 3.35 Å, corresponding to the (1 -2 -1) crystal plane of $rTiO_2$ (the phase is $Ti_9O_{17}$, 00-050-0791). The interplanar spacing of rMSN was 1.88 Å, which corresponded to the (5 2 1) crystal plane of MSN (P6/mmm, ICSD#48153). Notably, we observed that $rTiO_2$ may have been partially coated with some MSN crystals. This was confirmed by the analysis of lattice fringes of $rTiO_2$ and MSN, respectively. (**e**) Inverse fast Fourier transform (FFT), FFT, and crystal structures of $rTiO_2$ (top) and rMSN (bottom), respectively. Inverse FFT confirmed its stable phase, and the FFT image profiles were similar to the corresponding crystal structure, which further confirmed the stable phase even after high-temperature $N_2/H_2$ reduction. (**f**) HRTEM image and atomic ordering intensity in two different selected directions (top and bottom) of $rTiO_2$-MSN. The relatively stable atomic arrangement and interplanar spacing confirmed the stable crystalline phases.

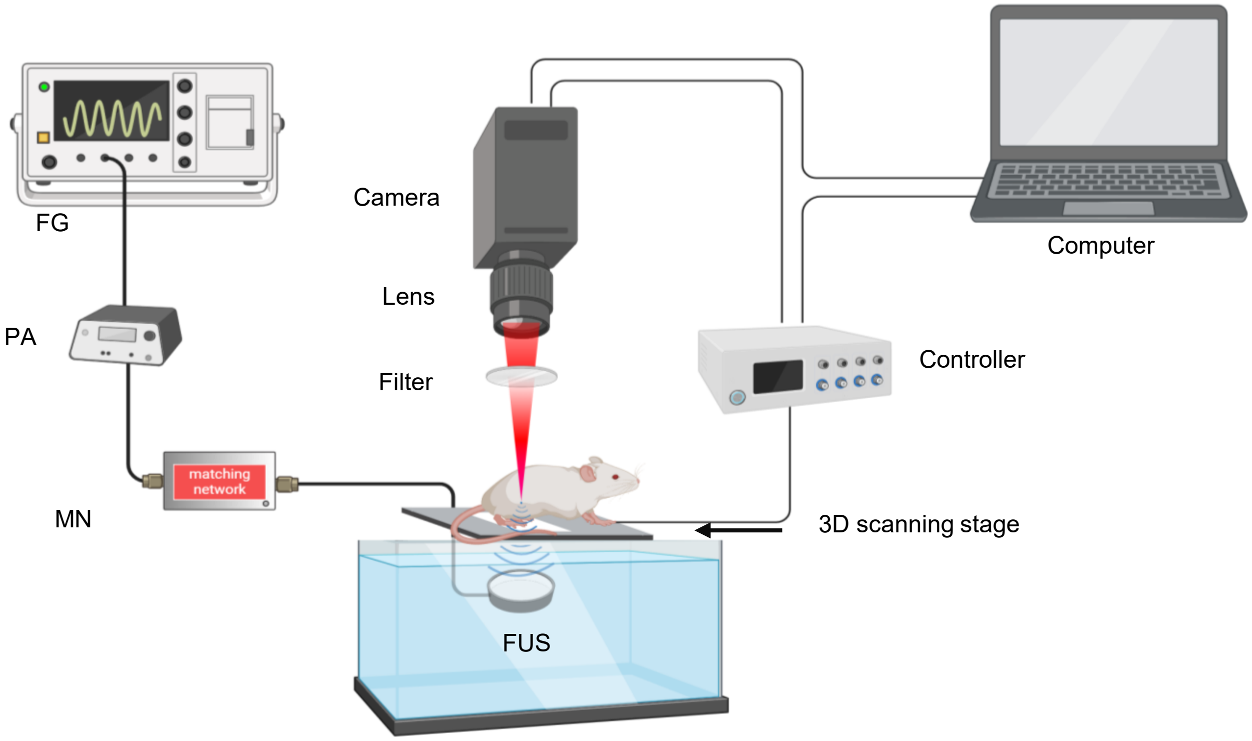


**Supplementary Figure 24.** Schematic diagram of the scanning focused ultrasound AL imaging system. Focused ultrasound was generated using an ultrasound transducer operating at ~4.55 MHz with a FWHM of ~1 mm. The ultrasound transducer was driven by a custom power amplifier with a sine wave input from a function generator (SDG2122X, SIGLENT), and impedance matching for the transducer was achieved using a matching network. The ultrasound-triggered AL was collected using a fixed-focal lens (SWIR-25, Navitar) and an InGaAs camera (C-RED 2 ER, Oxford Instruments). An NIR-II filter was used to filter the AL emission. A motorized stage (M-VP-25XL, Newport Corporation) was used for raster scanning and controlled by a LabVIEW program. The motorized stage and camera were synchronized via the stage controller (XPS-RLD4, Newport Corporation).

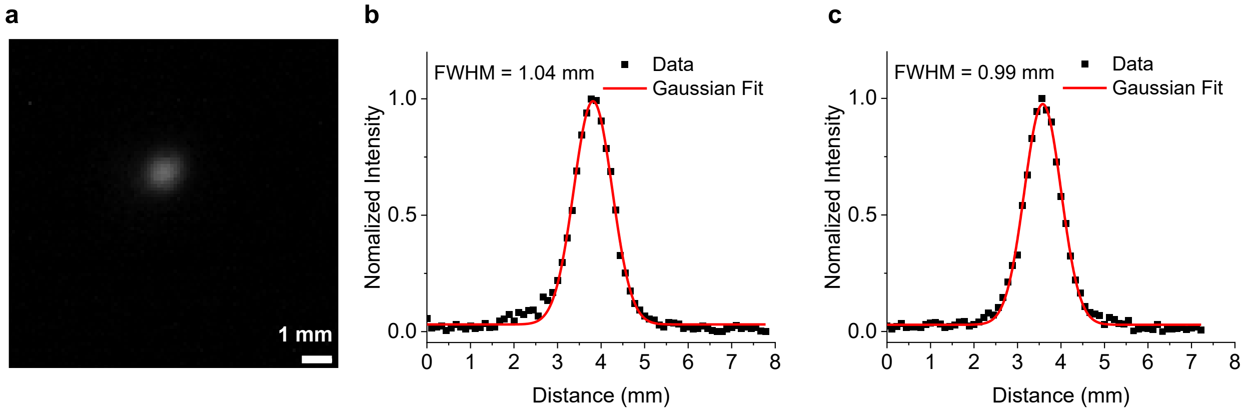


**Supplementary Figure 25.** The FWHM of the focus of the ultrasound transducer operating at ~4.55 MHz. To measure the FWHM, a ~0.5-mm-thick layer of powdered r$TiO_2$-MSN sample was embedded at the bottom of a PDMS substrate and positioned at the focused plane of the ultrasonic transducer. The AL signal at the focus was recorded by an InGaAs camera after being filtered by a 1500-nm long-pass filter. The FWHMs of the AL signal in the lateral and vertical directions were ~1.04 and ~0.99 mm, respectively.

**Supplementary Table 1. Comparison of current ultrasound-triggered luminescence materials**

| Materials | Wavelength (nm) | US frequency (MHz) | US power density ($W/cm^2$) | Ref. |
|---|---|---|---|---|
| $ZnS:Ag^{+},Co^{2+}$ | 470 | 1.5 | 10 | 6 |
| $CaZnOS:Er^{3+}$ | 500 - 600, 627 - 700 | 0.5 | 55.87 - 1117.4 | 7 |
| $CaZnOS:Nd^{3+}$ | 850 - 1000 | 0.5 | 1.4 - 141.5 | 8 |
| $CaZnOS:Mn^{2+}$ | 610 | 0.04 | 4 | 9 |
| $ZnGa_2O_4$: $Cr^{3+}$ | ~720 | 0.5 | 1.4 - 35.2 | 10 |
| Trianthracene derivative-based nanoparticles | ~600 - 750 (TD) | 0.03 | 4.5 - 9.2 | 11 |
| | | 1 | 1.5 - 2 | |
| Dioxetane-functionalized PDMS | 420 | 0.55 | 204 - 376 | 12 |
| **TMOs and REOs** | **> 1800** | **1** | **0.1 - 2.2** | **This work** |
